# Machine learning for characterizing plant-insect interactions through electrical penetration graphic signal

**DOI:** 10.1101/2024.06.10.598170

**Authors:** Quang Dung Dinh, Daniel Kunk, Truong Son Hy, Nalam Vamsi, Phuong D. Dao

**Affiliations:** Institut Galilée, Universite Sorbonne Paris Nord, Villetaneuse 93430, Paris, France; Department of Agricultural Biology, Colorado State University, Fort Collins, CO 80523, United States; Department of Cell and Molecular Biology, Colorado State University, Fort Collins, CO 80523, United States; Graduate Degree Program in Ecology, Colorado State University, Fort Collins, CO 80523, United States; Department of Mathematics and Computer Science, Indiana State University, Terre Haute, IN 47809, United States

## Abstract

The electrical penetration graph (EPG) is a well-known technique that provides insights into the feeding behavior of insects with piercing-sucking mouthparts, mostly hemipterans. Since its inception in the 1960s, EPG has become indispensable in studying plant-insect interactions, revealing critical information about host plant selection, plant resistance, virus transmission, and responses to environmental factors. By integrating the plant and insect into an electrical circuit, EPG allows researchers to identify specific feeding behaviors based on distinct waveform patterns associated with activities within plant tissues. However, the traditional manual analysis of EPG waveform data is time-consuming and labor-intensive, limiting research throughput.

This study presents a novel machine-learning approach to automate the segmentation and classification of EPG signals. We rigorously evaluated six diverse machine learning models, including neural networks, tree-based models, and logistic regressions, using an extensive dataset from aphid feeding experiments. Our results demonstrate that a Residual Network (ResNet) architecture achieved the highest overall waveform classification accuracy of 96.8% and highest segmentation overlap rate of 84.4%, highlighting the potential of machine learning for accurate and efficient EPG analysis. This automated approach promises to accelerate research in this field significantly and has the potential to be generalized to other insect species and experimental settings. Our findings underscore the value of applying advanced computational techniques to complex biological datasets, paving the way for a more comprehensive understanding of insect-plant interactions and their broader ecological implications. The source code for all experiments conducted within this study is publicly available at https://github.com/HySonLab/ML4Insects.

**Author summary:** Insect pests of the order Hemiptera pose a significant threat to global agriculture, causing substantial crop losses due to direct feeding and serving as vectors for many economically important plant viruses. Understanding plant-insect interactions is crucial for mitigating these impacts. The electrical penetration graph (EPG) is a valuable tool that provides detailed insights into these interactions. However, the analysis of EPG data is a time-consuming, labor-intensive process that can also be prone to operator errors. State-of-the-art machine learning (ML) algorithms can be trained to perform this task accurately and consistently. These advanced algorithms can automate identifying and classifying specific EPG waveform patterns associated with distinct insect feeding behaviors. Our machine learning models, trained on extensive aphid feeding data demonstrated high accuracy in classifying these waveforms, with Residual Network (ResNet) architecture achieving the best performance. The automated approach saves time and resources, eliminates operator error, and also enables the identification of novel feeding patterns, providing a deeper understanding of the mechanisms underlying plant-aphid interactions. Moreover, our evaluation of a large, diverse dataset of four aphid species on four host plants indicates the potential for generalizing these models to different experimental settings. By applying advanced computational techniques to EPG data, we are pioneering the intelligent surveillance of aphid feeding habits. This approach promises to significantly enhance our efforts in developing a better understanding of factors that affect aphid feeding.

## Introduction

Aphids (order Hemiptera) are major agricultural pests that feed on plant phloem sap using their needle-like mouthparts. Their rapid asexual reproduction and high population densities can lead to significant crop damage. Copious feeding by aphids not only depletes essential nutrients from plants but also introduces toxic saliva and plant viruses, further compromising crop quality and yield. In severe cases, aphid infestations can result in hundreds of millions of dollars in economic losses, underscoring the importance of understanding and managing these pests [1]. The electrical penetration graph (EPG) technique, introduced in 1964 by Donald McLean and Marvin Kinsey, provides researchers with a tool to develop a better understanding of aphid feeding behavior. [2]. Prior to EPG, real-time observation of aphid feeding was limited to histological methods, which only provide snapshots of the feeding process. The initial EPG technique involved creating an electrical circuit by wiring the aphid and the plant, followed by recording the resulting voltage changes once the aphid closes the circuit by penetrating the plant tissue. Throughout the 20th century, entomologists enhanced this technique by analyzing the behavioral mechanisms behind EPG waveforms using methods like stylectomy and histology. This allows us to uncover the underlying electrical component of each behavioral waveform, and through the invention of the direct current (DC) EPG system, which allows for the recording of behaviors originating from both plant-insect resistance (Ri) and those originating from the electromotive force (emf) [3, 4]. Today, the EPG technique is the predominant method for studying hemipteran feeding behavior, largely supplanting earlier histological methods.

Currently, commercially available DC-EPG systems provide the necessary hardware and software for acquiring EPG recordings [6]. To set up and create the circuit, aphids are restrained, and a gold wire is glued to their abdomen using conductive paint. The gold wire is typically soldered onto a brass nail, which can be inserted into a specially designed housing attached to the EPG circuit. For the plant, an electrode is inserted into the soil connected to the EPG circuit. The analog signal is converted to a digital signal in GIGA systems and recorded on a computer using the Stylet+ data acquisition software (.DAQ file and .DX extension for hour-by-hour recordings). After recording feeding behaviors, the user manually annotates the waveforms by visual inspection, creating a file with columns for the start time of the behavior, the specific behavior, and the voltage value at the start. Various feeding behaviors are then manually calculated from these annotation files or using publicly available Excel workbooks [7].

However, this method’s feasibility is greatly constrained because it demands a time-consuming manual annotation process and the expertise of an experienced observer to annotate the signal accurately, preventing high-throughput processing. During each annotation session, the observer analyzes data on a second-by-second basis, which can take up to 30 minutes, depending on the complexity of the observed patterns and the length of the recording. Thus, an automated label generation process will greatly streamline data analysis. Previous attempts at automation, such as Aphid Auto EPG [8], had limited classification ability due to its basic discrimination rules based on waveform amplitudes. Similarly, Adasme-Carreño et al. (2015) developed a system Assisted Analysis of EPG (A2EPG) to automate the annotation of EPG signals [9].

However, the criteria for distinguishing distinct waveforms was overly simplistic, leading to unsatisfactory predictions and the failure to identify or misclassify specific waveforms. Researchers have attempted to develop and implement machine learning methods for EPG signals due to their robustness to various data types and consistent predictive capabilities. Xing et al, (2023) proposed a classification model based on the Extreme Kernel Machine (EKM) to analyze the EPG signal of the pea aphid (*Acyrthosiphon pisum*) and the bird cherry-oat aphid (*Rhopalosiphum padi*), reporting a mean annotation rate of 94.47% [10]. Additionally, a Random Forest model was introduced to characterize the behaviors of Asian citrus psyllids (*Diaporina citri*) on nine citrus genotypes [11], achieving an overall accuracy score of 97.4%, with the Hidden Markov Model further revealing previously unknown feeding patterns. However, these studies were limited in scale, necessitating further validation of larger and more diverse datasets to assess the broader applicability of these methods. In recent years, Convolutional Neural Networks (CNNs) have shown great promise in insect behavior characterization [19–23], with their ability to automatically learn informative features from raw data eliminating the need for manual feature engineering. Furthermore, CNN variants have proven effective in segmenting insect biological time series [24–26]., showcasing their potential in EPG analysis.

In this study, we aim to 1) evaluate the performance of well-established machine learning models for annotating and characterizing EPG signals; 2) evaluate the effect of different feature extraction methods on ML classification model performance; and 3) develop a ready-to-use pipeline for automatic aphid feeding behavior classification for users not familiar with ML tools. Our study builds on previous work by evaluating various ML models, including *Fully Convolutional Network* (1DCNN), *2D Convolutional Neural Network* (2DCNN), *Residual Network* (ResNet), *Extreme Gradient Boosting Machine* (XGB), *Random Forest* (RF), and *Logistic Regression* (LR). A Python package was developed based on well-established libraries such as Scikit-learn and PyTorch, providing an end-to-end automatic classification process for easy reproducibility and widespread use.

## Materials and methods

### Data

The machine learning models were benchmarked on datasets collected from four aphid species: *Rhopalosiphum padi* (bird cherry-oat aphid; BCOA1, BCOA2) *Myzus persicae* (green peach aphid; GPA), *Phorodon cannabis* (cannabis aphid; CA) and *Aphis glycines* (Soybean aphid; SA) that were obtained from the *Stylet+* application [6]. The datasets include a set of signal data in ASCII format and a text file that provides corresponding ground-truth annotations. Each EPG recording consists of a single signal channel that spans 8 hours at a sampling rate of 100Hz. Waveforms within the EPG signals are classified into seven categories: non-penetrating (NP), pathway (C), potential drop (pd), phloem salivation (E1), phloem-feeding (E2), xylem feeding (G), and derailed stylet phase (F), depending on the pattern observed from both the time and frequency domain Fig 1. During the NP phase, the insect has yet to insert its mouthpart into the host plant, so the waveform is usually almost a straight line. Pathway phase (C) is when the stylet is inserted into the plant tissue and is present in the epidermal and mesophyll tissues. During C, brief intracellular punctures by stylet tips occur, referred to as pd or potential drops. It is during C that aphids make decisions regarding host acceptance or rejection. The E1 and E2 phases are sometimes combined and called the sieve element phase (SEP). During SEP, the stylet tips are within phloem sieve elements. During E1, the aphid actively salivates while maintaining stylet position within the sieve elements. The E2 phase signifies active phloem sap ingestion, with the stylet remaining in the sieve element. In contrast, the G waveform indicates xylem sap ingestion, characterized by the stylet positioned in the xylem. Lastly, the F waveform represents derailed stylets, where the aphid encounters mechanical difficulties while probing host tissue. In this phase, despite being within host tissue, the aphid does not engage in feeding behaviors due to stylet malfunctions. It is important to note that these behaviors are sequential. For instance, the aphid must enter C before engaging in phloem or xylem feeding (E1, E2, or G), and the pd pattern is only observed during C. Examples of waveforms of these behaviors are shown in Fig 1. Fig 2 describes the components of each dataset.

**Fig 1.**
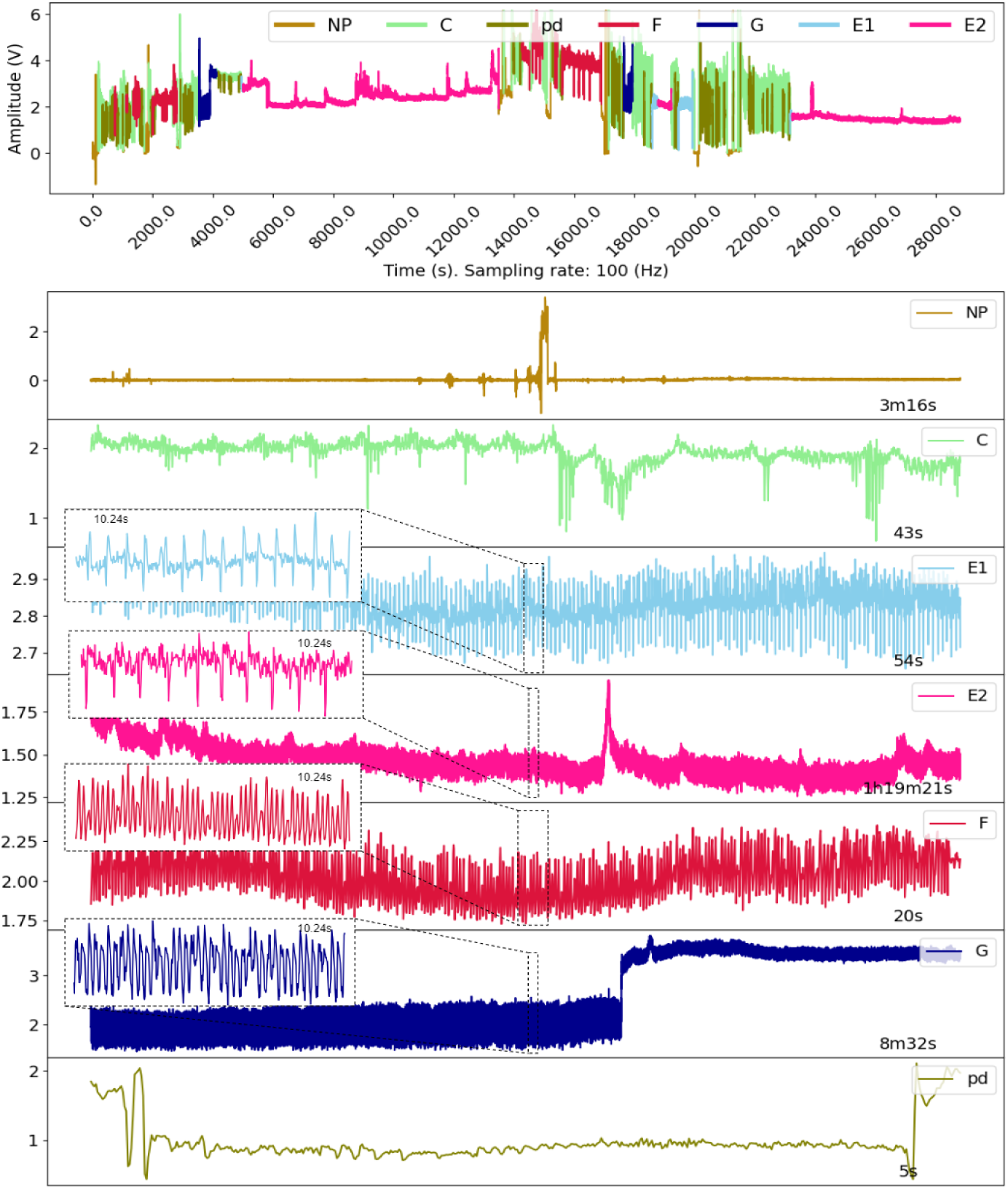
Representative waveforms observed during aphid feeding. The top panel is an example of a full EPG recording over 8 h or 28,800 s. The following panels show the segmented waveforms of seven different waveforms or behaviors observed during aphid feeding. In the NP or non-probing phase, the waveform is often almost flat, around 0, but occasionally has noisy peaks. Meanwhile, the pathway phase (C) labels include a complex range of behaviors, and the pd phase occurs within C. The sieve element phase consists of E1 and E2, and the derailed stylet (F) and xylem phase (G) are very distinctive in terms of their shapes and frequencies/periods. It should be noted that waveforms of these behaviors can vary depending on the aphid species and host plant(s), as well as the environmental conditions of the experiment.

**Fig 2.**
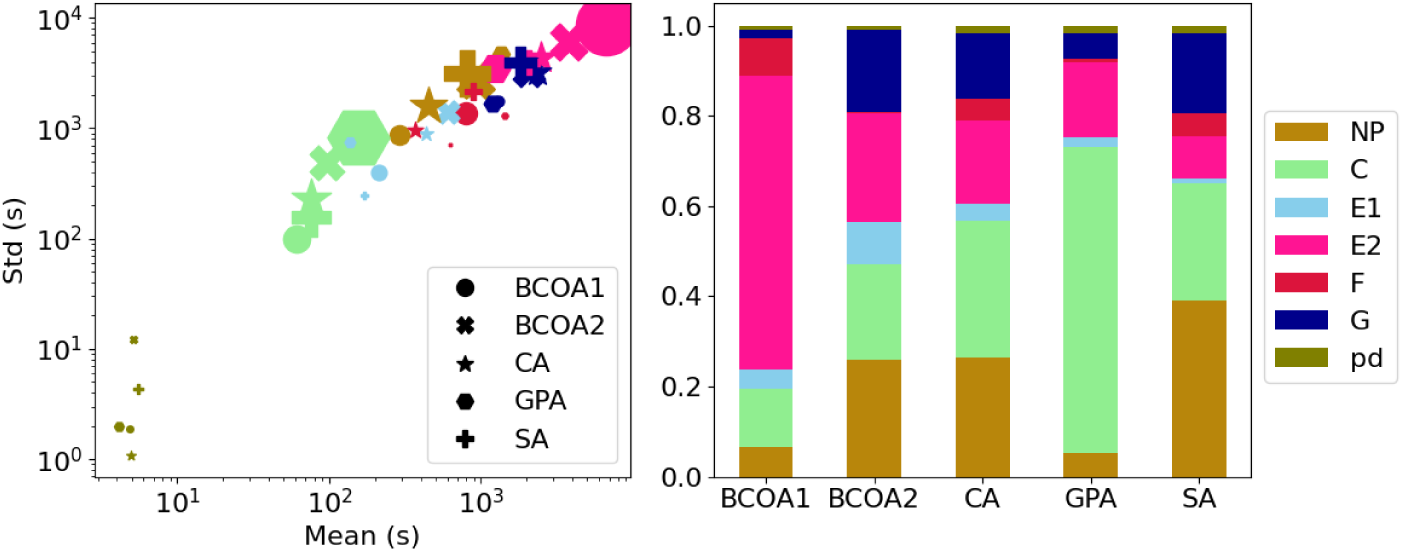
Details of the tested EPG dataset. The label distribution (left) versus the mean and standard deviation of each label (right). The size of each marker is proportional to the ratio of the corresponding label in the dataset.

Typically, the patterns from the pathwave phase (C) and sieve element phase (E1, E2) account for a major proportion of the total recording time. In two datasets, GPA and BCOA2, there were virtually no F and G waveforms, which partly explains the difficulty in distinguishing between these waveforms and the others. The potential drop (pd) waveform tends to appear frequently but over a short time, so they account for a negligible proportion of the data distribution. Next, we provide detailed meta-data for each experimental dataset in Table 1.

**Table 1.** Description of the studied datasets.

| Dataset | n | Species | Host plants | Host variety |
| --- | --- | --- | --- | --- |
| BCOA1 [12] | 121 | Bird cherry-Oat aphid | *Triticum aestivum* | Chinese Spring - susceptible |
| BCOA2 [13] | 57 | Bird cherry-Oat aphid | *Triticum aestivum* | Coker 9553 (AgriPro®) - susceptible |
| CA [14] | 62 | Cannabis aphid | *Cannabis sativa*<br>*Solanum tuberosum* | Elite<br>CO07015-4RU |
| SA [15] | 32 | Soybean aphid | *Glycine max* | PI 567301B - resistant<br>Williams 82 - susceptible |
| GPA [16] | 60 | Green Peach aphid | *Arabidopsis thaliana* | Col-0 (wild type)<br>*atg5* and *atg7* (autophagy mutants) |

**BCOA1** [12]. This dataset is derived from an experiment on the bird cherry-oat aphid (*R. padi*) feeding on wheat (*Triticum aestivum* var. Chinese Spring). Aphid feeding was recorded for 8 h over a 24-hour period during the morning, afternoon, and night under two different light conditions: a photoperiod of 16 h light / 8 h dark and continuous 24-hour light. The aim of the experiment is to understand how the time of day and light-dark cycles impact feeding behavior. Aphids were allowed to feed from the 2nd leaf of seedlings at Zadoks stage Z1.2 (reference to be added). The characteristics of their feeding waveforms exhibited minimal variation across the different experimental conditions.

**BCOA2** [13]. This dataset is derived from an experiment conducted on the bird cherry-oat aphid (*R. padi*) feeding on Wheat (*Triticum aestivum* var. Coker 9553, AgriPro@) . Aphid feeding was recorded at the beginning of the day and the beginning of the night to understand the influence of the time of day on aphid feeding behavior. Aphids and experimental plants were reared in a 16 h light and 8 h dark photoperiod at 24 ± 1 °C. Experiments were conducted on seedlings at Zadoks stage Z1.2.

**CA** [14]. This dataset comes from an experiment conducted on the cannabis aphid (*P. cannabis*) feeding either on its primary host, Hemp (*Cannabis sativa* var. Elite, New West Genetics) or a secondary host, potato (*Solanum tuberosum* var. CO07015-4RU). The aim of this experiment was to understand the differences in the feeding behavior of viruliferous and non-viruliferous aphids on hemp and potato to determine the potential of the cannabis aphid to vector potato virus Y to these species. Experimental plants were reared in a greenhouse in a 16 h light / 8 h dark photoperiod at 21 ± 1°C / 16 ± 1°C, and two three-week-old hemp and potato plants were used in the experiments. Aphids performed poorly on potatoes, probably due to the difference in host plant physiology and cellular anatomy, and waveform characteristics between aphids feeding on potatoes and hemp plants are likely not identical.

**SA** [15]. This dataset is from the soybean aphid (*A. glycines*) feeding on two genotypes of *Glycine max* (soybean). The feeding behavior of aphids on the susceptible cultivar, Williams 82, was compared to the resistant plant introduction, PI 567301B, which contains the *Rag5* gene, on both whole plants and detached leaves to determine the origin of Rag5 mediated resistance. Experimental plants and aphids were reared in a 16 h light / 8 h dark photoperiod with a temperature of 24 ± 1°C, and plants at the V1 stage were used for all recordings. As experiments were conducted on whole plants and also detached leaves, waveform characteristics across treatments are likely not identical.

**GPA** [16]. This dataset is from experiments with green peach aphid (*M. persicae*) feeding on Arabidopsis. The aim is to understand the relationship between plant autophagy and insect feeding behavior and performance. Aphid colonies and experimental plants were reared in a 16 h light and 8 h dark photoperiod at 22 ± 3 °C. Aphids were allowed to feed on one-month-old wildtype Col-0 plants or autophagy mutants *atg5.1* and *atg7.2*. Aphids showed reduced feeding on mutant *atg7.2* ; however, waveform characteristics remained consistent across all experimental treatments.

## Data processing

### Preprocessing

#### Data splitting

For each dataset, we used 90% of the number of recordings (Subset 1) for training and classification evaluation (Task 1) and reserved the other 10% (Subset 2) for segmentation (Task 2) evaluation. The detail of the two evaluation tasks are discussed in the results section. Subset 1 recordings were divided into fixed-length segments in the initial segmentation stage. By default, we set this length to be 1024 time steps or 10.24s. This set of segments is then split into a training set (70%), a validation set (10%), and a test set (20%). The details of data splitting for training and Task 1 evaluation are given in Table 2.

**Table 2.**
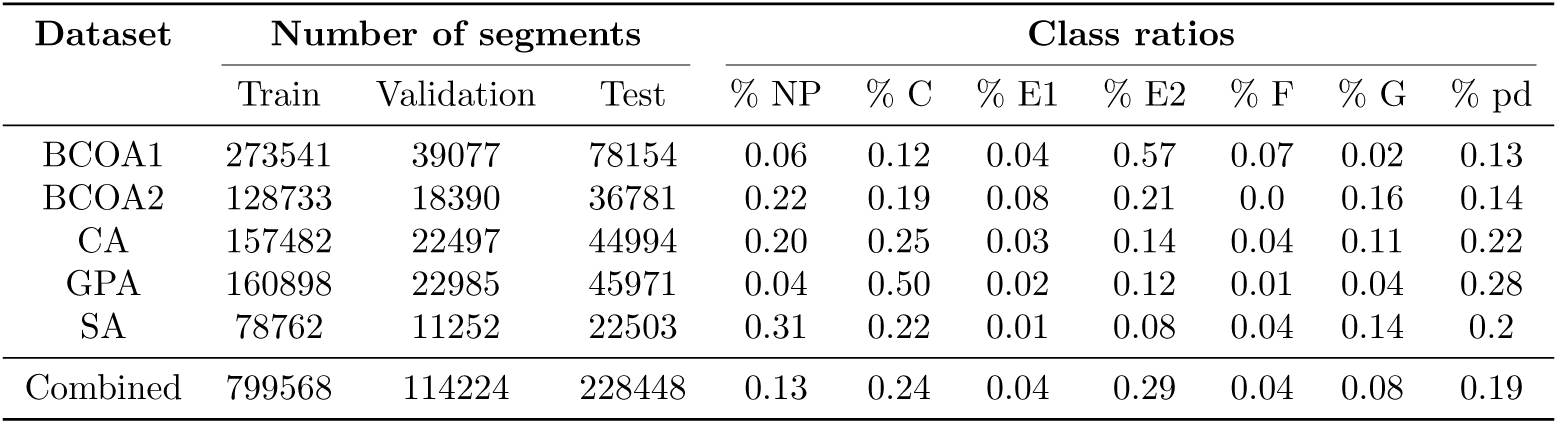
Data splitting for training ML models. Splits of five individual datasets and a combined dataset for training, validation, and evaluating the classification performance of the ML models in Task 1. For each dataset, recordings were broken down into segments, shuffled, and then split into training, validation, and testing subsets. The combined dataset, a concatenation of all five individual datasets, was also split similarly for further analysis and benchmarking.

Next, we describe the data preprocessing techniques. In the following, we denote by **x** *∈* R*^L×d^* a signal of *L* time steps and *d* channels.

#### Normalization

Data scaling is not mandatory but usually helps prevent numerical instability in calculations. We normalized each entire input recording **x** so that the amplitude ranges over [0, 1] by the formula

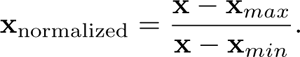

#### Augmentation

Recall that in the initial segmentation stage, we cut the input into fixed-length segments *w* (10.24s or 1024 time steps by default). Waveforms, of which the lengths are shorter than the predetermined value *w*, were extended to both sides.

Specifically, if the original waveform is *x* = (*x_i_, …, x_j_*), the padded waveform is given by

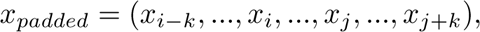

where 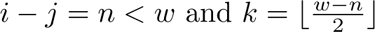

### Features extraction

*Fourier transform (FT)* (See Appendix. Fig S1): A technique that converts signals from the time domain (amplitude/time representation) to the frequency domain (amplitude/frequency representation) by assuming that the input signal is a linear combination of sine and cosine waves at different frequencies. FT identifies which frequencies are present in the input signal by determining the coefficients of these component waves. This method has proven to be highly effective in many time series classification frameworks because it captures the characteristics of time series data well in their frequency domain representation [30–33]. We fixed the hop length at 14 and the window size at 128 for generating spectrogram features.

#### *Wavelet transform (WT)* (See Appendix. Fig S2)

A powerful technique in signal processing that assumes the input signal is a linear combination of a family of special functions called *wavelets*, which are obtained by translating and scaling a *mother wavelet*. For our study, the *Morlet* wavelet family with geometric scales of length 64 and the Symlet4 wavelet family with 3 levels of decomposition were used for the continuous WT and discrete WT, respectively. The two versions work based on the same principle, but we shall discuss the discrete version only for simplicity. The discrete WT decomposes the original signal into several phases. In each phase, the signal is transformed into two complementary sets of wavelet coefficients: *high frequency* (detail, or ”D”) and *low frequency* (approximation, or ”A”) components. This is achieved by translating a wavelet along the time axis and convolving it with the input signal. The input signal is initially decomposed into ”A1” and ”D1”. In each subsequent phase, the *mother wavelet* is scaled by a factor of 2, and the same decomposition process is applied to the approximation coefficients from the previous stage using this scaled wavelet. Wavelet transform has proven effective in various signal processing tasks, including analyzing human biological signals [40, 42].

#### *Gramian Angular Field (GAF)* See Appendix. Fig S3

A technique that convert an time-series into image format by representing the rescaling time-series, transform it into polar coordinates, and finally represent it as a matrix comprises of the trigonometric sum (Gramian Angular Summation Field - GASF) or difference (Gramian Angular Difference Field - GADF). This representation allows us to apply 2D convolutional networks to time-series classification. In our study, GASF is used.

#### Statistical features

Mean, standard deviation, skewness, and quantiles are examples of these features, offering insights into data distribution. These features can capture the central tendency, dispersion, shape, and outliers of the data distribution, making them valuable for identifying changes or anomalies. Essentially, we can view these calculations as a method for dimensionality reduction because they can describe the data through a much smaller set of numbers. The specific statistical features used in this study include mean, root mean square, standard deviation, variance, skewness, the quantiles at level 0.05, 0.25, 0.5, 0.75, 0.95, and the zero crossing rates of the consecutive segments.

#### The complexity measures

Shannon entropy and permutation entropy provide valuable insights into a time series’s regularity, long-range autocorrelation, and order structure. Permutation Entropy (PE) is a particularly robust tool for analyzing time series, as it quantifies the complexity of a dynamic system by examining the order relations between values and deriving a probability distribution of ordinal patterns. We use permutation and Shannon entropy because they are straightforward, quick to compute, and provide substantial information. The two entropy measurements and the statistical features are either calculated on the raw signal, the Fourier coefficients, or the discrete WT coefficients.

### EPG signal annotation and characterization

Our EPG signal characterization approach consists of two main processes: dividing the signal into a set of consecutive fixed-length segments using a *sliding-window*, as detailed in [10, 11], followed by annotating each segment using classification algorithms. The task can be divided into three main steps: initial segmentation, prediction, and aggregation, as shown in Fig 3. The input signal is initially divided into consecutive segments of fixed-length *d*. The next step focuses on transforming the obtained segments using Fourier or Wavelet transformation and extracting valuable features from the transformed data. The Data processing section discusses this step in more detail. The ML models are then trained to learn from the transformed data to output a probability distribution, assigning the label of the highest probability to each segment. Each segment yields *d* predicted labels, which will be concatenated to create a complete segmentation of the input recording.

**Fig 3.**
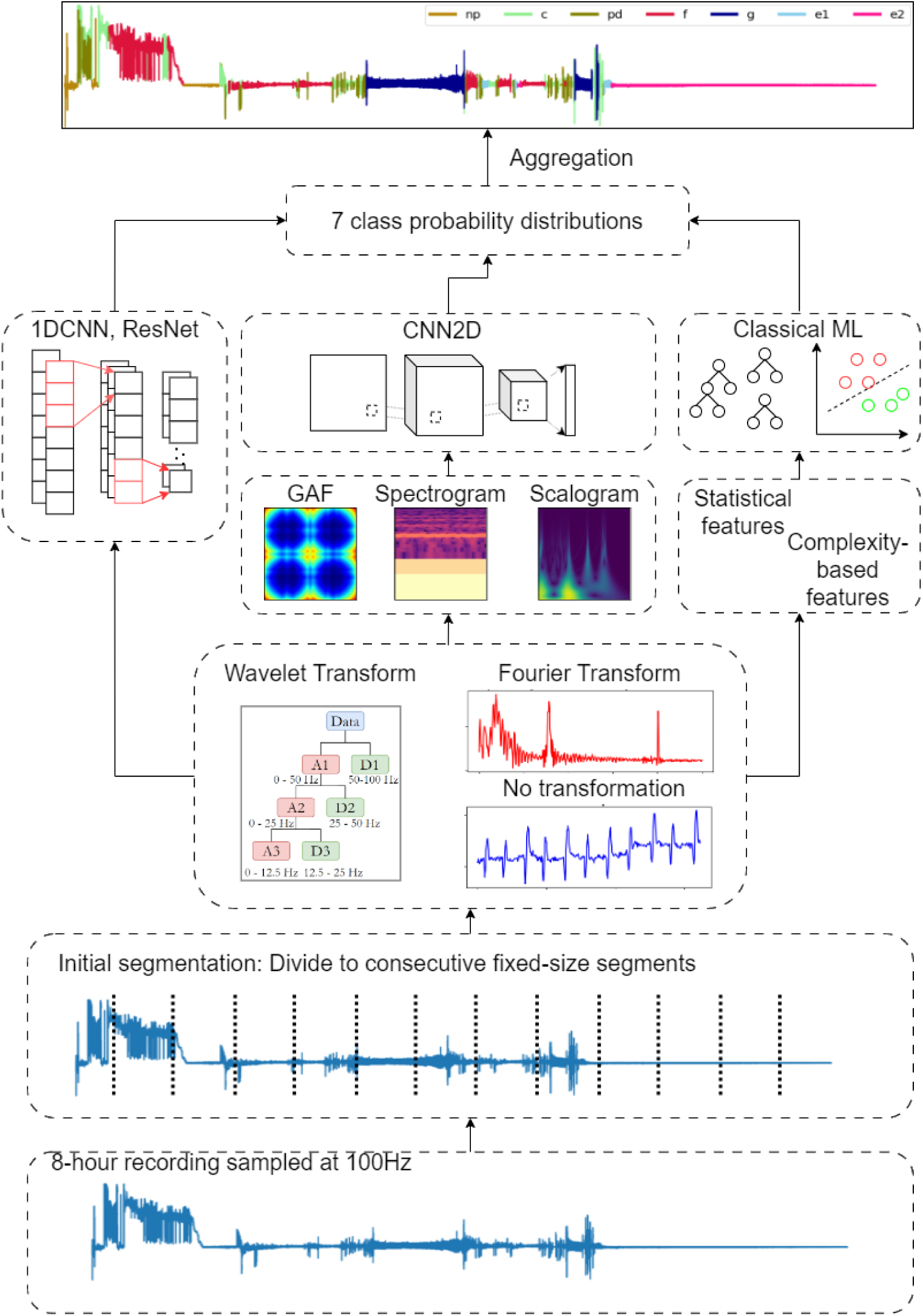
Automated EPG Signal Annotation and Characterization Pipeline. The figure illustrates the end-to-end process of automating EPG signal analysis. The input recording is divided into fixed-length segments, which are then passed to the features extraction stage. ML models learn the extracted features or raw signals to predict the behavior. Each segment is assigned to a class of the highest probability. Finally, the predicted labels are concatenated to form a complete annotated waveform.

### Non-deep learning classification algorithms

#### Random Forest (RF)

A well-known machine learning algorithm used for both classification and regression tasks. An RF consists of multiple Decision Trees (DTs) trained on subdatasets created by bootstrapping from random subsets of features. The time series adaptation of random forest [28] [29] involves segmenting the input time series into intervals and extracting various features from these segments. Experimental studies show that TSF using simple features such as mean, standard deviation, and slope is computationally efficient and outperforms strong competitors such as 1-NN classifiers with dynamic time-warping distance.

#### Extreme gradient boosting (XGB)

A very fast and accurate state-of-the-art tree-based models built under the Gradient Boosting framework. The algorithm operates by sequentially adding weak learners to the ensemble, with each new learner focusing on correcting the errors made by the existing ones while using a gradient descent optimization technique to minimize a predefined loss function during training. In [39], three state-of-the-art gradient boosting-based algorithms, including XGBoost, LGBM, GBM, and CatBoost, are benchmarked based on 12 datasets of different sizes and contexts. The results show that all models achieved state-of-the-art performance, with LGBM ranking first. However, it is crucial to note that extensive fine-tuning was essential to attain this level of performance.

#### Logistic Regression (LogReg)

A popular and widely used classification algorithm owing to its simplicity, interpretability, and effectiveness for binary classification tasks. Moreover, this method can be extended to multi-class classification problems through one-vs-all or softmax regression methods.

### Deep learning classification algorithms

#### *Convolutional neural networks (CNNs)* (See Appendix. Fig S4)

A type of neural network that learns characteristics of the input data by optimizing the *kernels* and *filters*. In CNNs, learning units are also arranged in layers, and the heart of CNNs is the *matrix convolution* operator. There are 3 types of layers in CNNs: convolutional, pooling, and fully-connected. At each convolutional layer, the input convolves with the kernels in order to produce *feature maps*. In other words, the kernel allows the network to concentrate on specific parts of the input sequentially, thereby enabling CNNs to minimize the parameters per layer and promote a more extensive network architecture. The pooling layer reduces the overall complexity of the model but in a trade-off for the loss of information during training. Finally, the feature maps are flattened into a single-dimensional array before being passed through fully connected layers to derive classification probabilities.

### Model accuracy assessment

The performance of the ML models is assessed based on their capability to classify individual input segments (Task 1) and their robustness in predicting insect behavior across the entire waveform (Task 2). The latter evaluation task involves comparing the overlap between the concatenated predicted waveform and the manually annotated waveform, which reflects the extent to which the predicted signal agrees with the ground-truth signal. To measure classification results, we utilize the accuracy and F1 scores calculated using the following equation.

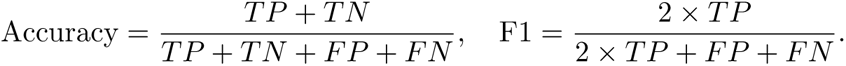

More specifically, overall accuracy (OA), the per-class accuracy, and the average F1 score (avg-F1) are calculated from predictions on the test set. To evaluate the robustness of the models, we measure the overlap and offset between the predicted and the manually annotated waveform. Recall that to produce a complete segmentation, one first segments the recording into consecutive segments of a size of 1024 time steps (10.24s) and classify each segment. Then, we give each time step (e.g., each 0.01 second) a predicted label according to that of the segment containing it; consequently, each segment yields 1024 labels for 1024 time steps. A complete predicted waveform is thus composed by concatenating these labels. Finally, we evaluate the prediction accuracy by calculating the percentage of agreement between the two versions of dense labelings (the *overlap rate*, or OR). For example, for an 8 h recording, we calculate the overlap between the 2880000 predicted labels and 2880000 ground-truth labels. The evaluations are done using a 10-fold cross-validation scheme to avoid biases, and the mean of these metrics is reported.

### Experimental settings

The 1DCNN model consists of three convolutional layers that output 64, 128, and 64 channels, respectively. Each layer uses a dilation of 3 and a stride of 2, followed by ReLU activation and batch normalization. After these convolutional layers, there is a max pooling layer with a size of 3. The feature maps are then flattened, subjected to dropout with a probability of 0.5, and passed through a fully-connected layer.

The ResNet model adopts the idea of skip-connection. This model comprises three skip-connection blocks, each containing three consecutive convolutional layers with *d*_1_, *d*_2_, and *d*_3_ filters, respectively, followed by ReLU activation and a skip connection, and finally, normalization. We set the number of output channels for each block to be 64, 128, and 128. After each convolution block, max pooling with a kernel size of 3 and a stride of 3 is performed to reduce computational complexity. The outputs of the convolutional layers will be flattened out, and a dropout with a probability of 0.5 will be applied for normalization before being passed through a final fully connected layer.

The 2DCNN model includes two convolutional layers, each outputting 16 channels and using a 3 *×* 3 kernel with a stride of 1. Each convolutional layer is followed by ReLU activation and max pooling with a size of 2 and a stride of 2. The results are then flattened and passed through two fully-connected layers with dropout to obtain the probability distribution for the 7 classes. The dropout rates are set to 0.5 for both fully-connected layers, with kernel size and stride for the max pooling layers set to 2.

For tuning of deep learning algorithms, the baseline hyperparameters are set to *{*256, 0.0001, 9, 100*}*, representing the batch size, learning rate, kernel size, and number of training epochs, respectively. To find the optimal set of parameters for an 1DCNN, these values are selected from the following ranges: batch size *{*128, 256, 512, 1024, 4096, 8192*}*, learning rate *{*0.01, 0.001, 0.0001, 0.00001*}*, kernel size *{*5, 9, 15*}*, and number of training epochs *{*20, 50, 100*}*. We start by fixing the learning rate and kernel size, then manually select the batch size from *{*128, 256, 512, 1024, 4096, 8192*}*. We then fix the batch size at 256 and experiment with learning rates of *{*0.01, 0.001, 0.0001, 0.00001*}*, achieving the highest mean validation accuracy (mVA) of 0.902 with a learning rate of 0.0001. The same procedure is applied to the kernel size and the number of training epochs. The mVA is used to determine the best set of hyperparameters. Regarding the loss function, we use the Cross-Entropy loss function, which is typical for multilabel classification tasks.

For traditional ML algorithms, we configured XGB with 100 trees (*n* = 100), a learning rate of 0.3 (*η* = 0.3), and a maximum tree depth of 6 (*d_max_*= 6). For Random Forest, *n* is set to 100, but *d_max_* is not specified. A grid search was conducted on XGB with the number of trees, learning rate, and maximum tree depth drawn from *{*50, 100, 200, 300*}*, *{*0.01, 0.1, 0.2, 0.3*}* and *{*3, 4, 5, 6*}*, respectively. The optimal hyperparameters found are *{*300, 0.3, 6*}*. For Logistic Regression, we used the default parameters provided by *scikit-learn*.

Our experiments were conducted on a personal laptop GIGABYTE G7 equipped with a 12-core Intel i5-10500H CPU (2.5GHz) processor and an NVIDIA GeForce RTX 3060 Laptop GPU (6GB memory). Training times ranged from half an hour to two hours, depending on the data volume, while the segmentation task was typically completed in a matter of minutes. All scripts of the deep learning architecture are implemented using the Pytorch framework [18], while the traditional machine learning models are implemented using *scikit-learn*. For signal processing tasks, we use *librosa* and *PyWavelet* to perform Fourier transform and wavelet transform, respectively.

## Results

For each ML model, we performed 3 groups of experiments with respect to 3 different techniques to preprocess its inputs, namely no transformation, Fourier transform, and wavelet transform (See Table 3). Each group contained experiments on 5 individual datasets having different settings. To assess the transferability of the ML models into different scenarios, we additionally conducted experiments with two representative models from the non-deep learning and deep learning groups.

**Table 3.** Input Structures of Machine Learning Models for EPG Signal Classification. Details of input structures to ML models with respect to the experimented signal processing and feature extraction techniques. In the table, *b* is the batch size, *d* is the predetermined length of each segment, and *n* is the number of training observations. The abbreviations of the feature columns are None: No further feature extraction, GAF: Gramian Angular Field, Spec: spectrogram, Scalo: scalogram, and Handcrafted: statistical and complexity features. For 2DCNN, all generated images are rescaled to either 64 *×* 64 or 65 *×* 65 according to the transformation technique.

| Model | Transformation | Feature | Input shape |
| --- | --- | --- | --- |
| 1DCNN, ResNet | Raw | None | $(b, 1, d)$ |
| | FT | None | $(b, 1, d)$ |
| | WT | None | $(b, 1, d)$ |
| 2DCNN | Raw | GASF | $(b, 1, 64, 64)$ |
| | FT | Spec | $(b, 1, 65, 65)$ |
| | WT | Scalo | $(b, 1, 64, 64)$ |
| LogReg, XGB, RF | Raw | Handcrafted | $(n, 13)$ |
| | FT | Handcrafted | $(n, 13)$ |
| | WT | Handcrafted | $(n, 52)$ |

**Table 4.** Performance of Deep Learning Models. Classification and segmentation performance of 1DCNN, ResNet, and 2DCNN on 5 datasets with respect to different signal feature engineering techniques. The bold figures highlight the best mean metric of a model.

| Dataset |  | 1DCNN |  |  | ResNet |  |  | 2DCNN |  |  |
| --- | --- | --- | --- | --- | --- | --- | --- | --- | --- | --- |
|  |  | Task 1 |  | Task 2 | Task 1 |  | Task 2 | Task 1 |  | Task 2 |
|  |  | OA | Avg-F1 | OR | OA | Avg-F1 | OR | OA | Avg-F1 | OR |
| Raw | BCOA1 | 0.928 | 0.862 | 0.870 | 0.968 | 0.962 | 0.948 | 0.775 | 0.531 | 0.751 |
|  | BCOA2 | 0.914 | 0.893 | 0.647 | 0.936 | 0.907 | 0.784 | 0.688 | 0.589 | 0.532 |
|  | CA | 0.925 | 0.920 | 0.857 | 0.956 | 0.951 | 0.869 | 0.670 | 0.594 | 0.478 |
|  | SA | 0.915 | 0.839 | 0.770 | 0.914 | 0.830 | 0.769 | 0.702 | 0.528 | 0.576 |
|  | GPA | 0.904 | 0.832 | 0.732 | 0.943 | 0.894 | 0.851 | 0.840 | 0.573 | 0.704 |
|  | Mean | **0.917** | **0.869** | **0.775** | **0.943** | **0.909** | **0.844** | 0.735 | 0.563 | 0.608 |
| FT | BCOA1 | 0.916 | 0.885 | 0.913 | 0.939 | 0.914 | 0.922 | 0.947 | 0.915 | 0.860 |
|  | BCOA2 | 0.895 | 0.877 | 0.740 | 0.911 | 0.904 | 0.742 | 0.919 | 0.887 | 0.748 |
|  | CA | 0.876 | 0.870 | 0.823 | 0.922 | 0.917 | 0.847 | 0.921 | 0.916 | 0.721 |
|  | SA | 0.892 | 0.822 | 0.725 | 0.913 | 0.863 | 0.718 | 0.923 | 0.842 | 0.649 |
|  | GPA | 0.871 | 0.838 | 0.838 | 0.888 | 0.821 | 0.840 | 0.913 | 0.847 | 0.805 |
|  | Mean | 0.890 | 0.858 | 0.808 | 0.915 | 0.884 | 0.814 | 0.925 | 0.881 | 0.757 |
| WT | BCOA1 | 0.938 | 0.907 | 0.918 | 0.953 | 0.929 | 0.941 | 0.954 | 0.935 | 0.933 |
|  | BCOA2 | 0.848 | 0.795 | 0.648 | 0.922 | 0.884 | 0.702 | 0.923 | 0.896 | 0.771 |
|  | CA | 0.904 | 0.878 | 0.826 | 0.928 | 0.917 | 0.865 | 0.921 | 0.914 | 0.869 |
|  | SA | 0.910 | 0.809 | 0.692 | 0.941 | 0.876 | 0.729 | 0.913 | 0.808 | 0.755 |
|  | GPA | 0.883 | 0.754 | 0.836 | 0.927 | 0.871 | 0.855 | 0.925 | 0.876 | 0.855 |
|  | Mean | 0.897 | 0.829 | 0.784 | 0.934 | 0.895 | 0.818 | **0.927** | **0.886** | **0.837** |

### Training on individual datasets

#### Deep learning

The 1DCNN and ResNet models excelled and yielded superior outcomes when trained with different types of inputs, showing the strong learning capability of these models. These models achieved near-perfect OA and average F1 scores across the three groups of experiments. The highest OAs for ResNet was 96.8%, 93.9%, and 95.3% with raw inputs, Fourier-transformed inputs, and wavelet-transformed inputs, respectively.

Likewise, those for 1DCNN were 92.8%, 91.6% and 93.8%, respectively. On average, the OAs of ResNet were 2.6%, 2.5%, and 3.7% higher than those of 1DCNN, while ResNet also demonstrates significantly better performance in terms of the avg-F1 scores, with 4.00%, 2.6% and 6.6% higher on average. ResNet worked noticeably well with simple raw input, achieving an average of 84.5% OR over the 5 datasets, with the highest accuracy of 94.8% observed in the BCOA1 dataset. However, using transformed inputs with FT and WT reduces the average OR of ResNet to 81.4% and 81.8%, respectively. This means that raw data were more informative to this model, as it is robust to capture relevant features from this input type. In fact, the waveforms of aphids are characterized only by certain frequency ranges, as mentioned by W.F. Tjallingii in [5]. Therefore, using all Fourier coefficients for training the model might be redundant. In contrast, this additional information was useful for 1DCNN, of which architecture is much simpler than ResNet, as it increases the mean OR of 1DCNN to 80.8% using FT inputs. In Fig 5, we visualize the training process of ResNet, 1DCNN, and 2DCNN through the loss and accuracy curve when training with raw inputs. A similar pattern can be observed when training with the other two types of inputs. The training process of 1DCNN experienced a *spiking* phenomenon, a common phenomenon in stochastic gradient descent, which should only raise concern if training or validation accuracy declines. Conversely, ResNet exhibited a much smoother loss curve and a tendency to overfit rapidly, especially after just 20 epochs. This underscores the importance of robust regularization techniques to enhance its generalization capabilities across diverse datasets. The per-class accuracy scores given by the confusion matrix in Fig 4 reveal that ResNet was slightly less efficient than that of 1DCNN when trained on a small dataset such as SA, indicating potential overfitting in the model training.

**Fig 4.**
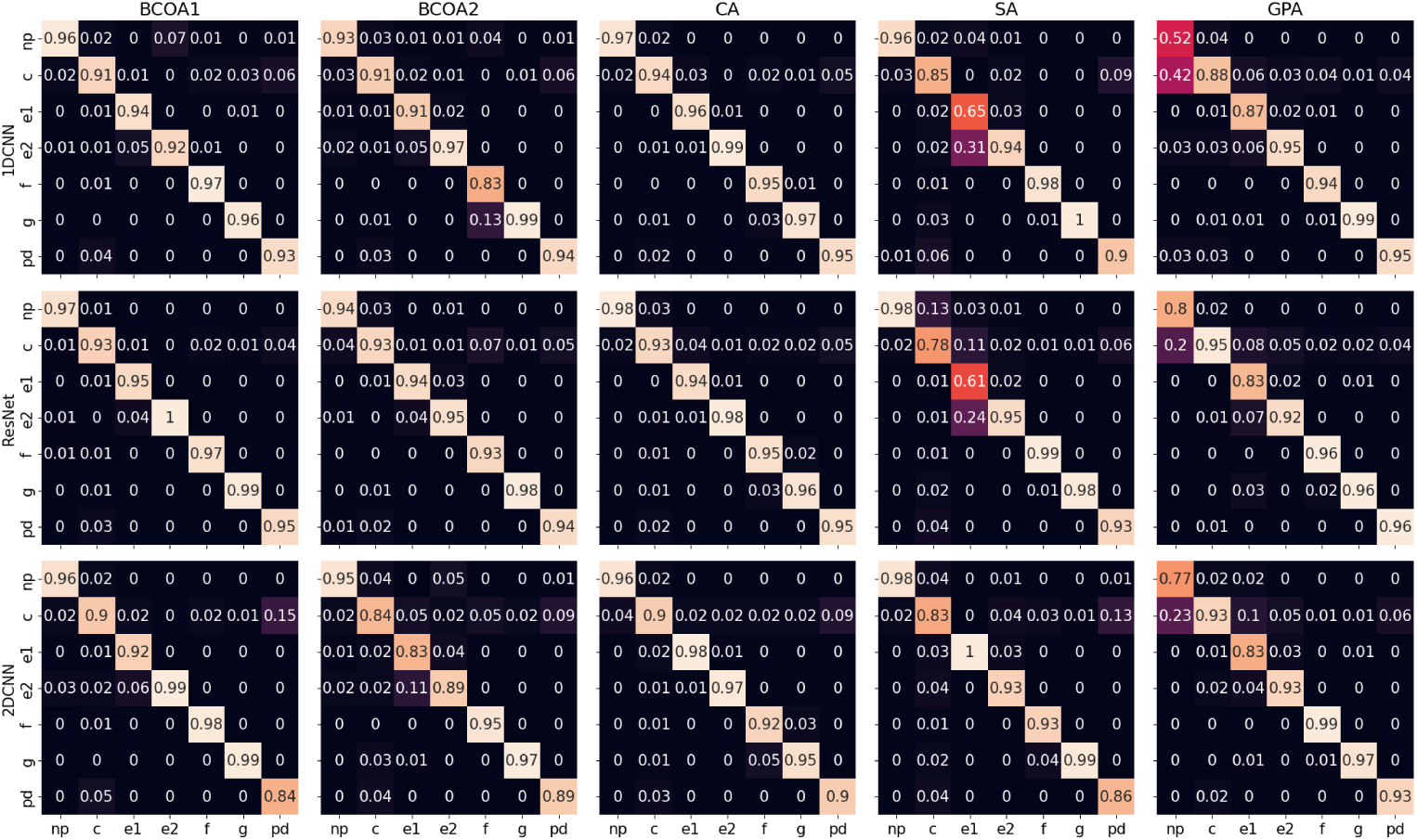
Confusion Matrices of Deep Learning Models for EPG Waveform Classification. The figure presents the confusion matrices for the three deep learning models in Task 1: 1DCNN with raw inputs, ResNet with raw inputs, and 2DCNN with scalogram input features. The confusion matrices illustrate the models’ performance in classifying each of the seven waveform categories (NP, C, pd, E1, E2 G and F). The diagonal elemnets represent correct classifications, while off-diagnonal elements indicate misclassifications.

**Fig 5.**
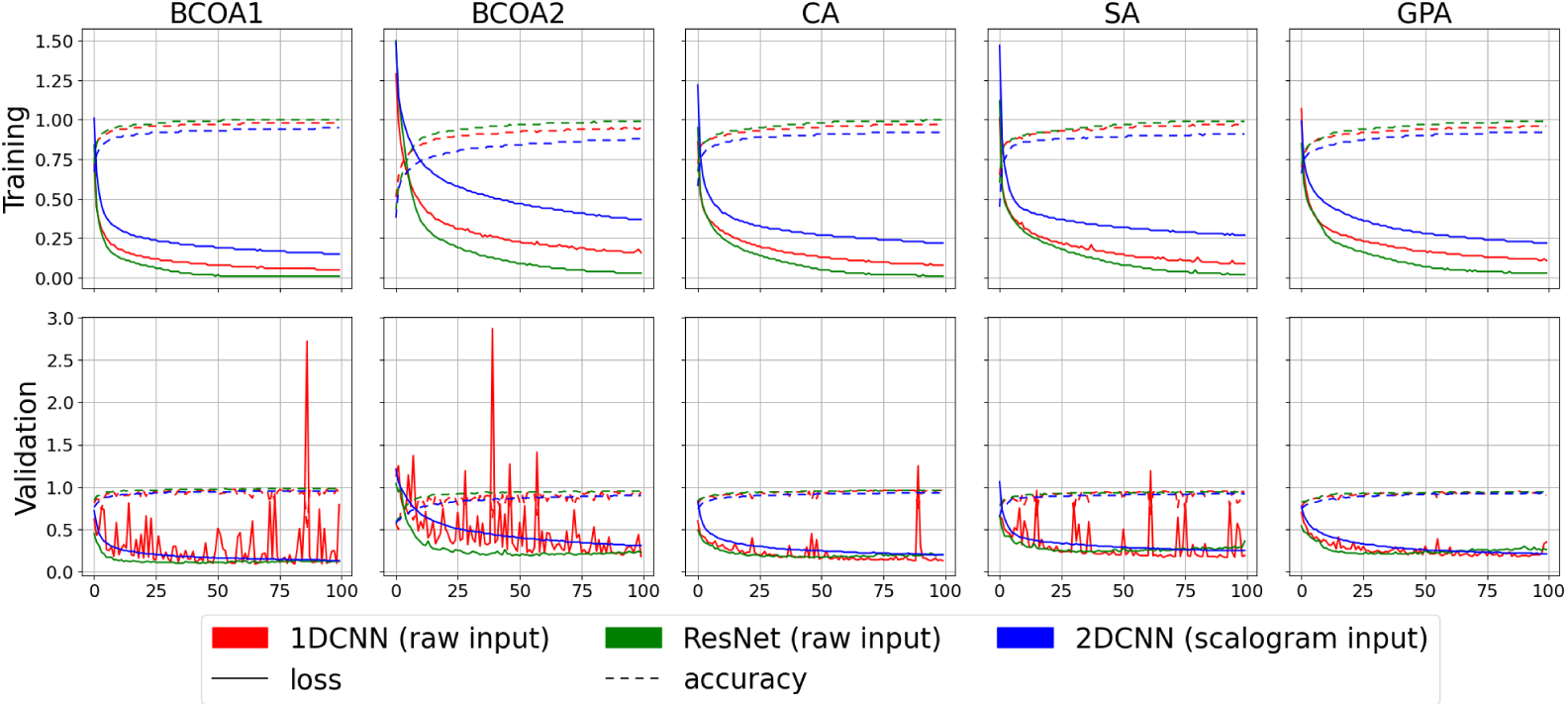
Training and Validation Accuracy and Loss Over 100 Epochs for Deep Learning Models. The graph illustrates the performance of three ML models (1DCNN, ResNet and 2DCNN) over 100 epochs of training and validation. The top and bottom panel displays the measurements for training phase and validation phase, respectively. Solid lines present the model loss (as measure of prediction error), and dotted lines present classification accuracy (proportion of correct predictions). The distinct colos correspond to each model: red for 1DCNN, green for ResNet and blue for 2DCNN.

The 2DCNN model differs from 1DCNN and ResNet as it requires image inputs. To implement this method, we first generate input images of segments using three different techniques: GAF, spectrogram, and scalogram. The model yielded the best performance when trained with scalogram inputs, reaching the highest OA and avg-F1 of 95.4% and 93.5% with the BCOA1 dataset. While the spectrogram inputs may produce similar results in Task 1, model performance with scalogram as input could achieve better generalization as there was an 8% increase in mean OR, from 75.7% with spectrogram to 83.7% with scalogram. We further investigated the confusion matrices of the 2DCNN model with scalogram input, as this approach yields the best performance. On all five datasets, we observed that 2DCNN can misclassify between (pd) and (C) waveforms, which is understandable as (C) is a complex waveform and (pd) is a part of (C). The scalograms of (pd) may have captured redundant frequency information. With the SA dataset, 2DCNN exceptionally reached a perfect accuracy for (E1), while ResNet and 1DCNN struggled to identify this waveform. Despite experiencing fast learning after the initial 20 epochs, the actual test scores of 2DCNN show that it can generalize quite well even after 100 epochs of training.

#### Traditional machine learning

Regarding the non-deep learning models, the Extreme Gradient Boosting (XGB) Classifier performed fairly well and achieved results that were comparable to the best case of the DL algorithms, especially to the best metrics reported by ResNet, and outperformed Logistic Regression in all cases. Features extracted from WT at various resolutions demonstrated the effectiveness in characterizing the EPG waveforms, which was also the most effective of the three signal processing techniques for XGB. Indeed, WT provides the models with richer information about the frequency domain of the input signal, with a total of 4 resolutions instead of 1 in Fourier transform. Moreover, the transformation principle of WT helps localize the waveform behavior regarding its frequency, which is crucial for characterizing EPG waveforms as they are defined based on the observed patterns across a certain range of frequencies. For classification task, the OAs range from 94.2% to 96.4%, while the avg-F1 ranges from 90.9% to 95.8%.

Details of the classification results are reported in Table 5. As a result, the OR reported by XGB with WT features demonstrates a robust classification performance, with an average OR of 84.5% across 5 datasets. Meanwhile, Random Forest is also a robust model, and its accuracy metrics closely follow XGB’s in all experiments. The highest mean OA, avg-F1, and OR metrics obtained by RF when trained on 5 datasets are 93.4%, 89.5%, and 81.8%, respectively. This is unsurprising as RF and XGB are built on the same basis of aggregating the results from a set of weaker learners -decision trees. Besides, using RF with the sliding-window approachs was thoroughly examined in [28, 29], which suggests the method’s adaptability to time-series classification. Overall, the two models demonstrated impressive performance across global scores and individual per-class scores, often exceeding 90% with only a few exceptions that were slightly lower (See Fig 6).

**Fig 6.**
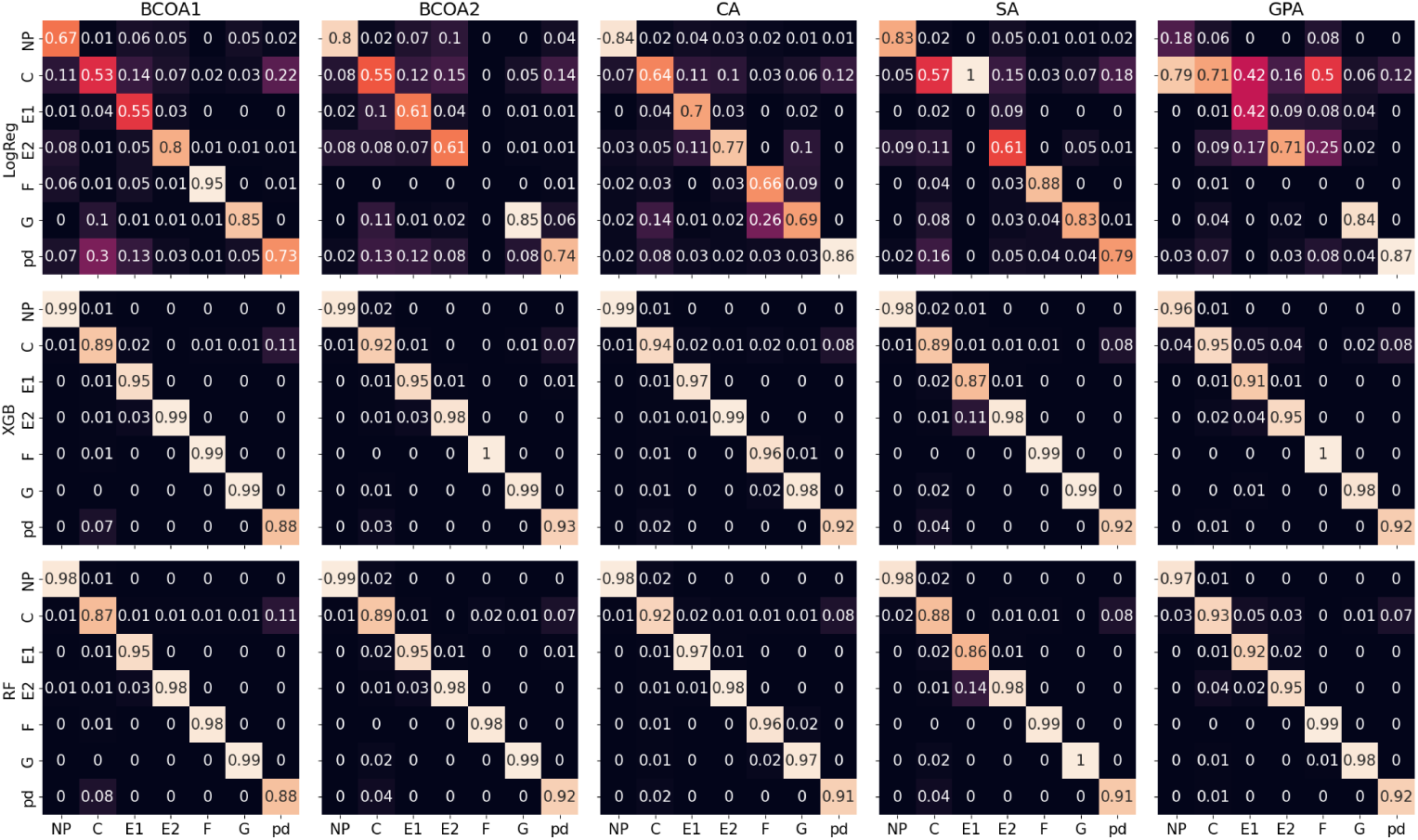
Confusion Matrices of Traditional Machine Learning Models for EPG Waveform Classification. The figure presents the confusion matrices for LogReg, XGB and RF models with features crafted from 3-level discete WT using *symlet4* mother wavelet. As before, the confusion matrices illustrate the models’ performance in classifying each of the seven waveform categories (NP, C, pd, E1, E2 G and F). The diagonal elemnets represent correct classifications, while off-diagnonal elements indicate misclassifications.

**Table 5.** Performance of Traditional ML models. Classification and segmentation performance of XGB, RF and LogReg on 5 datasets with respect to different signal feature engineering techniques. The bold figures highlight the best mean metric of a model.

| Dataset |  | XGB |  |  | RF |  |  | LG |  |  |
| --- | --- | --- | --- | --- | --- | --- | --- | --- | --- | --- |
|  |  | Task 1 |  | Task 2 | Task 1 |  | Task 2 | Task 1 |  | Task 2 |
|  |  | OA | Avg-F1 | OR | OA | Avg-F1 | OR | OA | Avg-F1 | OR |
| Raw | BCOA1 | 0.911 | 0.864 | 0.776 | 0.910 | 0.864 | 0.771 | 0.616 | 0.233 | 0.604 |
|  | BCOA2 | 0.945 | 0.939 | 0.602 | 0.947 | 0.942 | 0.589 | 0.541 | 0.401 | 0.555 |
|  | CA | 0.907 | 0.892 | 0.704 | 0.905 | 0.891 | 0.716 | 0.628 | 0.411 | 0.585 |
|  | SA | 0.915 | 0.865 | 0.631 | 0.914 | 0.870 | 0.628 | 0.706 | 0.286 | 0.644 |
|  | GPA | 0.879 | 0.778 | 0.798 | 0.878 | 0.779 | 0.794 | 0.534 | 0.281 | 0.497 |
|  | Mean | 0.881 | 0.867 | 0.702 | 0.911 | 0.869 | 0.699 | 0.605 | 0.322 | 0.577 |
| FT | BCOA1 | 0.886 | 0.822 | 0.841 | 0.879 | 0.813 | 0.833 | 0.673 | 0.244 | 0.653 |
|  | BCOA2 | 0.915 | 0.889 | 0.579 | 0.916 | 0.885 | 0.600 | 0.454 | 0.312 | 0.593 |
|  | CA | 0.907 | 0.892 | 0.705 | 0.878 | 0.862 | 0.705 | 0.587 | 0.378 | 0.580 |
|  | SA | 0.894 | 0.846 | 0.616 | 0.893 | 0.841 | 0.753 | 0.632 | 0.245 | 0.685 |
|  | GPA | 0.845 | 0.738 | 0.776 | 0.837 | 0.722 | 0.631 | 0.535 | 0.271 | 0.615 |
|  | Mean | 0.889 | 0.837 | 0.703 | 0.881 | 0.825 | 0.705 | 0.576 | 0.29 | 0.625 |
| WT | BCOA1 | 0.964 | 0.951 | 0.932 | 0.956 | 0.940 | 0.927 | 0.776 | 0.563 | 0.756 |
|  | BCOA2 | 0.962 | 0.955 | 0.789 | 0.955 | 0.952 | 0.778 | 0.691 | 0.577 | 0.709 |
|  | CA | 0.957 | 0.958 | 0.856 | 0.949 | 0.949 | 0.865 | 0.760 | 0.675 | 0.692 |
|  | SA | 0.951 | 0.909 | 0.787 | 0.945 | 0.892 | 0.750 | 0.757 | 0.573 | 0.73 |
|  | GPA | 0.942 | 0.927 | 0.863 | 0.931 | 0.905 | 0.852 | 0.751 | 0.400 | 0.73 |
|  | Mean | **0.955** | **0.940** | **0.845** | **0.947** | **0.928** | **0.835** | **0.747** | **0.558** | **0.723** |

### Training on the combined dataset

One of the main goals of our study is to develop a transferable model applicable to datasets derived from various aphid species, host plants, and experimental conditions. To enhance the analysis, we further evaluated the performance of two representative models, 1DCNN and XGB, on the combined dataset created by merging data from BCOA1, BCOA2, CA, SA and GPA. We select 1DCNN to train on raw data over the other candidates because of its computiationally simplicity and comparable performance compared to the other two methods. The XGB model with WT features was selected as it is the best model in terms of both training speed and performance metrics out of the non-deep learning models.

For XGB, the inputs include a total of 52 statistical and non-linear complexity features. Feature selection becomes pivotal to discern correlations and determine the features best suited for training. Employing Pearson’s correlation score, we identify features with correlations exceeding 0.8 as highly correlated. Fig 7 shows the correlations matrix before and after feature selection, in which we note that the feature group containing the k-quantiles *{*0.05, 0.25, 0.5, 0.75, 0.95*}* at all three WT detail resolutions are highly correlated, so we only keep the median (or the 0.5 quantile) features and discard the others. Moreover, the other feature groups consisting of standard deviation, variance, and root-mean-square also show strong correlations, as they are very similar in terms of calculation. We remove variance and root mean square features and only keep the standard deviation as representative for the three features for each resolution. The remaining 24 features include the means, standard deviations, skewnesses, zero-crossing rates, and the Shannon and permutation entropies of four resolutions A3, D3, D2, D1. This feature selection greatly improves the speed of the feature engineering and extraction step, alleviating the burdensome time requirements associated with these processes.

**Fig 7.**
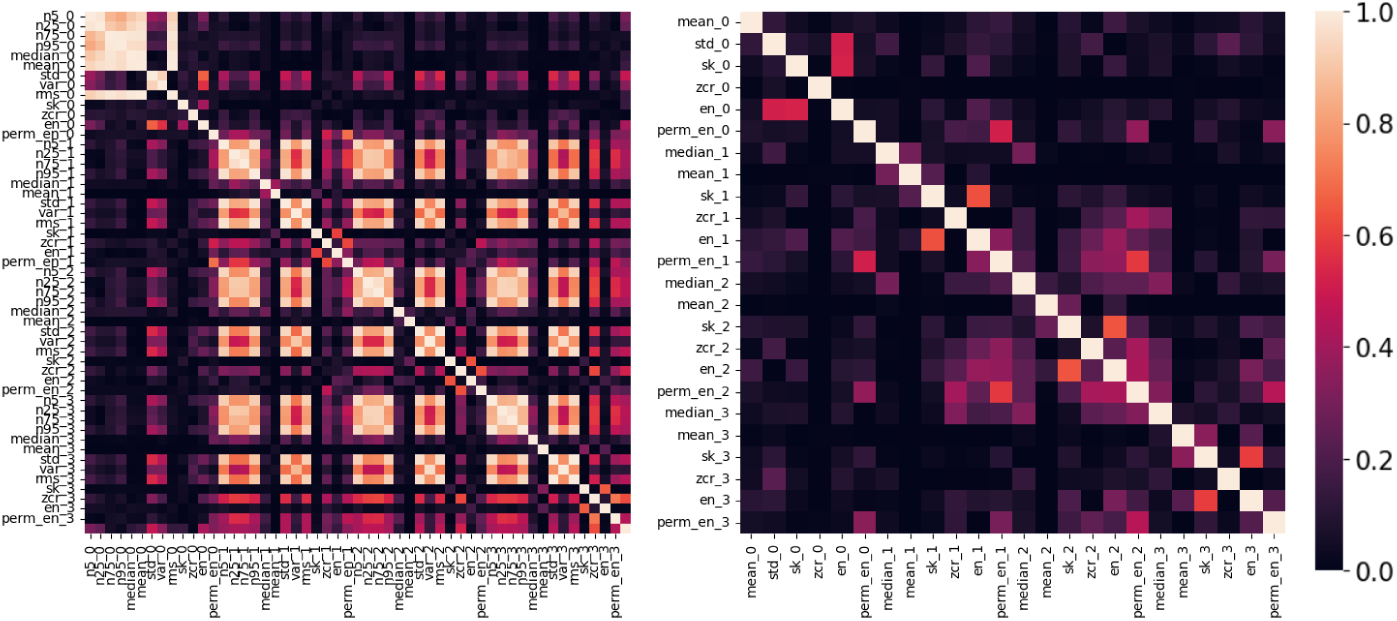
Correlation Matrices of Handcrafted Features Before and After Feature Selection. The figure presents two heatmaps illustrating the correlation among 52 handcrafted features derived from the 3-level wavelet transform (WT) at four different resolutions. These features were used as inputs for traditional machine learning algorithms. The left map demonstrates the correlation matrix of 52 features crafted from 4 resolutions of the 3-level WT, which are used as inputs for traditional machine learning algorithms. The right heat map is the correlation matrix after feature selections, in which features whose Pearson’s correlation coefficient is higher than 0.8 are considered highly correlated. We keep only one representative feature and remove its highly correlated features.

In addition, hyperparameter tuning for optimal parameter selection is critical in ML model training. We employed a Grid Search (GS) technique to fine-tune the XGB model. Initially, we conduct an exhaustive search to identify the best combination by manually exploring all possible scenarios. Table 6 shows the results of different training settings. Using a smaller set of features, XGB managed to attain a high classification accuracy for all the waveforms and only experienced a slight reduction in OA and avg-F1 metrics from 92% to 89.1% and from 92% to 88.9%, respectively. These results can be slightly improved by using the optimal set of hyperparameters obtained from Grid Search, which consequently yields 90.9% OA and 90.8% avg-F1. The most accurate prediction results were achieved using identical hyperparameters on the original dataset with 52 features.

**Table 6.** Features Selection and Hyperparameters Tuning Results of XGB. Per-class accuracies, overall accuracy, and macro-average F1 score of the XGB method on the combined dataset.

| Dataset | Per-class accuracy |  |  |  |  |  |  | OA | Avg-F1 | OR |
| --- | --- | --- | --- | --- | --- | --- | --- | --- | --- | --- |
|  | NP | C | E1 | E2 | F | G | pd |  |  |  |
| FS + GS | 0.954 | 0.844 | 0.870 | 0.956 | 0.948 | 0.960 | 0.870 | 0.909 | 0.908 | 0.808 |
| FS | 0.942 | 0.816 | 0.838 | 0.940 | 0.937 | 0.946 | 0.860 | 0.891 | 0.889 | 0.805 |
| GS | **0.978** | **0.888** | **0.924** | **0.975** | **0.972** | **0.976** | **0.888** | **0.936** | **0.937** | **0.841** |
| Baseline | 0.970 | 0.859 | 0.897 | 0.964 | 0.958 | 0.963 | 0.875 | 0.920 | 0.920 | 0.836 |

For 1DCNN, we employed a more aggressive strategy due to the computational complexity involved in the model training. This involves manually adjusting hyperparameters in a fixed sequence: batch size, learning rate, kernel size, and then the number of epochs. The effectiveness of each hyperparameter option was evaluated based on the mean validation accuracy (mVA). A total of 16 experiments were conducted, and the outcomes are detailed in Table 7. Ultimately, we determined that a batch size of 256, a learning rate of 0.0001, and a minimum kernel size of 9 yield the best outcomes. The left sub-figure of Fig 8 shows the training curves of the 1DCNN models with a batch size of 256, learning rate of 10^−4^, and kernel size of 9. As seen, a number of training epoch from 20 to 50 was generally sufficient for the model to achieve robust learning. The middle and right figures show the final metrics for both tasks of the 1DCNN models at an epoch of 100. The 1DCNN model achieve a high classification scores, at 92.1% OA and 92% avg-F1, respectively. The per-class accuracies showed that the highest misclassification is frequently observed between (C) and (pd) throughout our studies, primarily due to the mixture and overlap of the two behaviours. Meanwhile, the OR rates across the 5 proposed datasets were 85.5%, 81.5%, 65.3%, 83% and 63.1% for BCOA1, BCOA2, CA, GPA and SA, respectively. We observe low ORs in dataset that contains irregular samples of aphids’ behavior such as CA and SA, which indicates that the uniformness of recorded EPG signal can affect the overall quality of the ML models.

**Fig 8.**
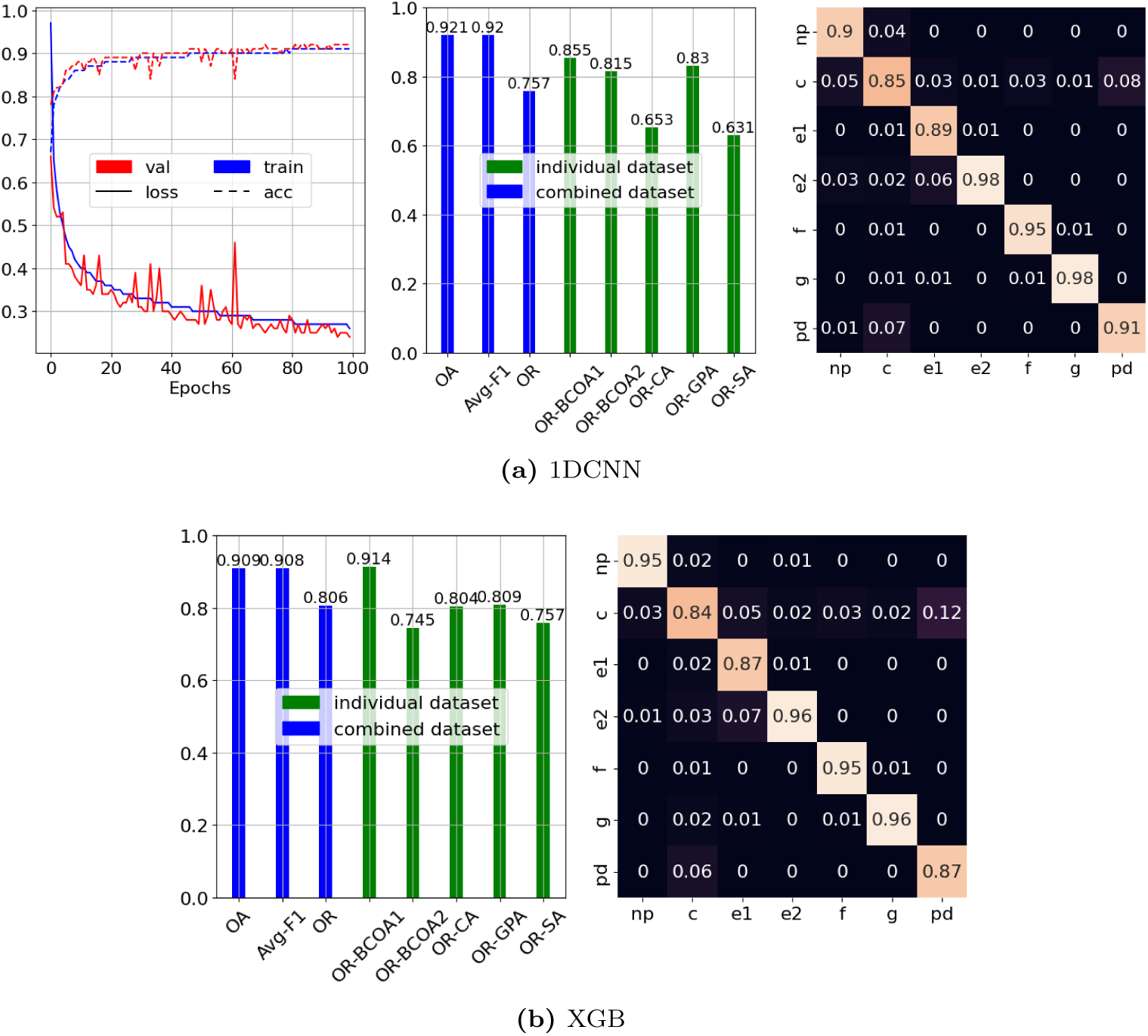
Performance of 1DCNN Models with Specific Parameters for EPG Signal Analysis. The figure illustrates the performance of 1DCNN models trained with a batch size of 256, a learning rate of 10^−4^, and a kernel size of 9 on EPG signal data. The left panel shows the loss (solid lines) and accuracy (dotted lines) curves during the training (blue) and validation (red) phases. These curves provide insights into the model’s learning progress and ability to generalize to unseen data. The middle panel presents evaluation metrics for both tasks at epoch 100. The Overall Accuracy (OA) is the proportion of correct predictions across all classes. The Average F1 Score (avg-F1) is a balanced measure of precision and recall, considering false positives and false negatives. The mean overlap rate (mean OR) is the average overlap between predicted and true waveform segments, measuring the accuracy of temporal segmentation. Each dataset’s overlap rate (OR) provides insights into dataset-specific performance variations. The right panel shows the confusion matrix for the classification task, revealing the per-class accuracy and any misclassifications between waveform categories. The diagonal elements represent correct classifications, while off-diagonal elements indicate errors. This comprehensive analysis provides a detailed view of the 1DCNN model’s performance, encompassing both the learning process and the final evaluation on multiple metrics and datasets.

**Table 7.**
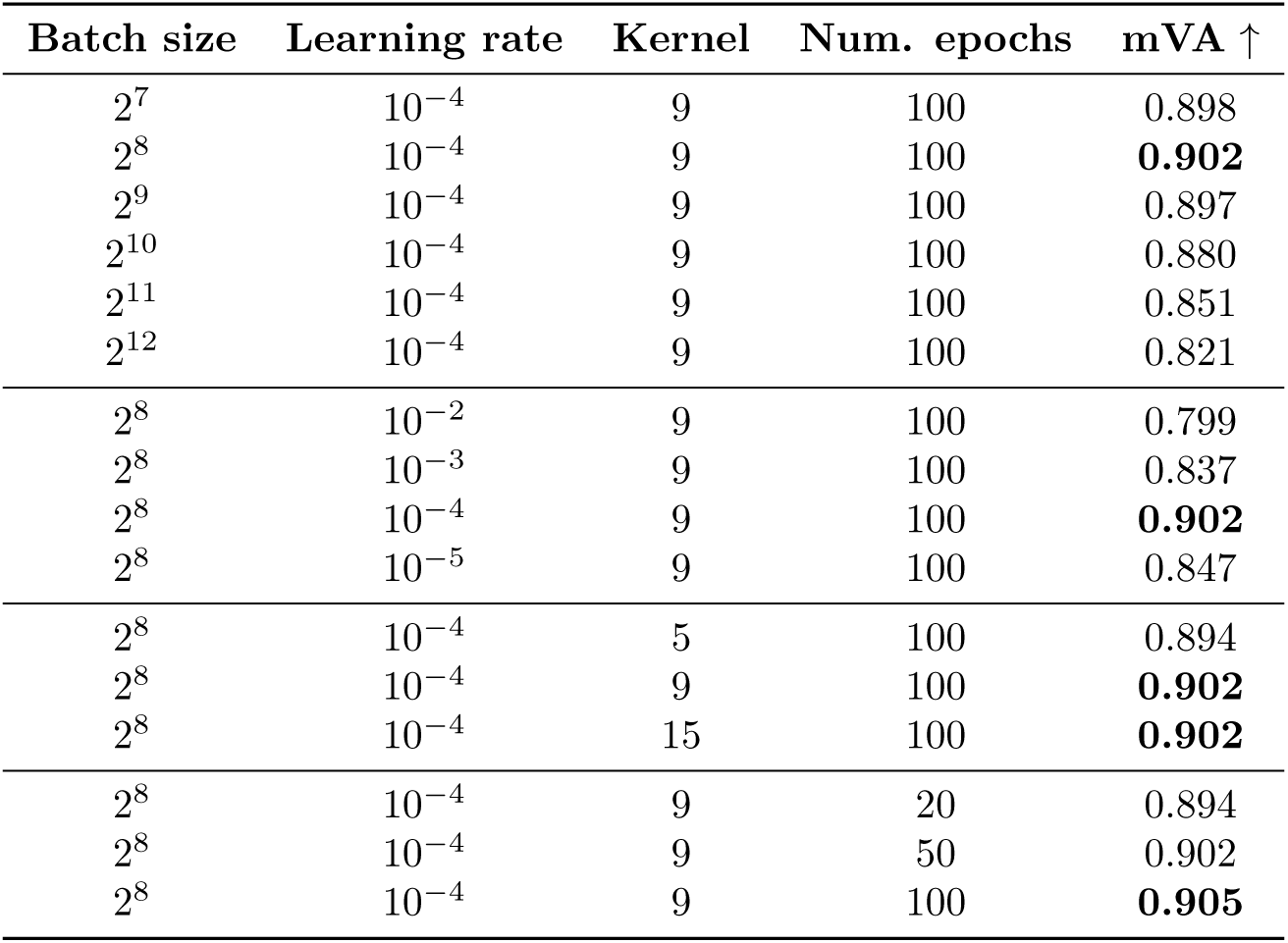
The Hyperparameters Tuning results of 1DCNN. The table shows the hyperparameters tuning results of 16 experiments for 1DCNN on the combined dataset. The best model was selected based on the mean Validation Accuracy (mVA). We found that *{*2^8^, 10^−4^, 9*}* was the best setting for batch size, learning rate and kernel size, while 20-50 training epochs usually ensures a good learning result.

## Discussion

This study presents the benchmark results of six machine learning models in EPG signal annotation for characterizing aphid feeding behavior. The annotation procedure was implemented through three main steps: initial segmentation, segment classification, and labeled segment aggregation. Each model was evaluated thoroughly based on two criteria: waveform classification (Task 1) and full annotated waveform quality assessment (Task 2). For Task 1, the prediction accuracy of individual segments was evaluated, including the classification accuracy calculation and the F1 score. Meanwhile, the overlap rate between the predicted and the ground-truth annotated waveforms was estimated for Task 2. Five aphid datasets collected from four aphid species and five different host plants were used to develop and test the ML models. The results show that ML models, in general, are robust enough to annotate and characterize EPG signals and aphid-feeding behavior. Two representative models (1DCNN and XGB) were selected based on their prediction accuracy and computational complexity to be additionally evaluated on a large aggregated dataset. Our generalized models can be transferred to other datasets collected from different experimental conditions and host plant species.

Next, we provide a more detail view on two situations where the automatically generated annotations have high and low OR. Fig 9 shows an example of the predicted annotation from the 1DCNN model that almost perfectly match the ground-truth version, reaching a commendable OR of 95%. Oftenly, long and dominant waveform such as G, F, E2 are correctly identified and located. Instances of non-overlap are primarily due to the offset between the predicted and ground-truth waveforms. This is an unavoidable issue with the sliding approach since we classify the time steps every consecutive 1024 time steps, e.g., from 0 to 1024, from 1024 to 2048, from 2048 to 3072, until we reach the end of the input recording. On the other hand, the natural behavior of aphids is not fixed in such a way, leading to this misalignment of the waveform endpoints. The phenomenon is particularly profound with short waveforms such as (pd), often overlooked by ML models. Thus, it is advisable for users to conduct a post-prediction evaluation to mitigate these minor inaccuracies.

**Fig 9.**
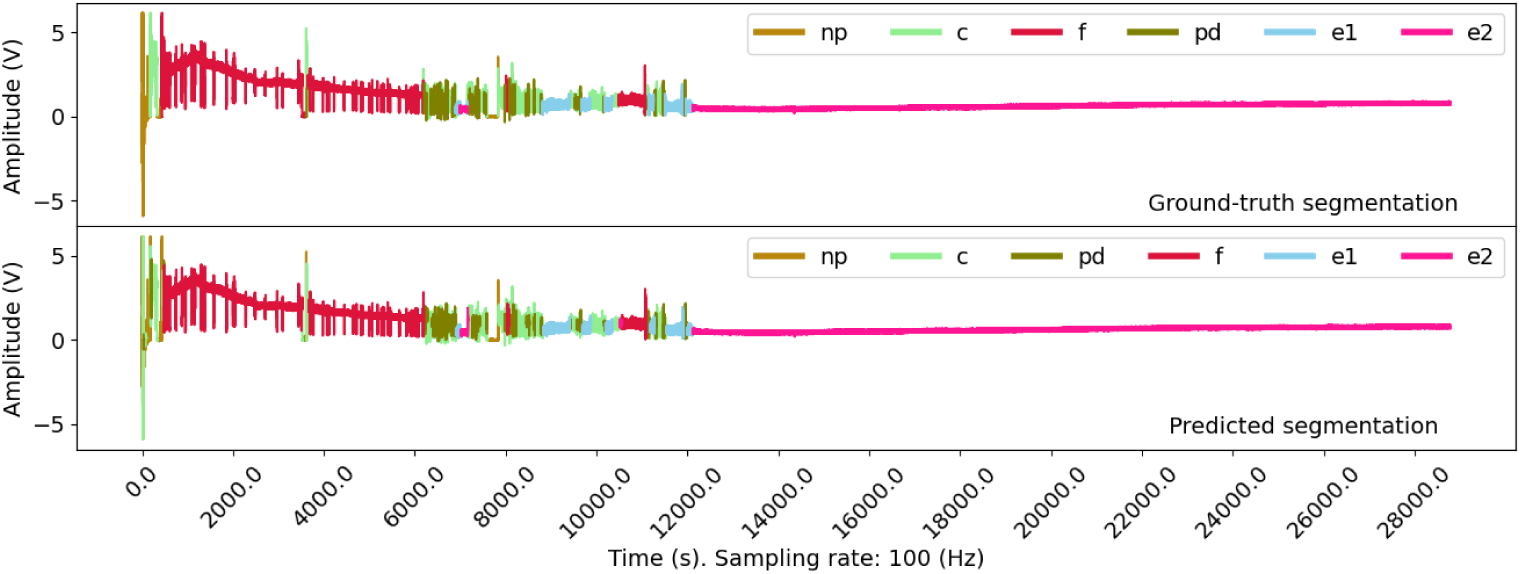
Prediction of aphid behaviours from input EPG recording. An example of ground-truth and model-predicted waveforms for a recording in the BCOA1 dataset. The behaviors of the experimented aphids are consistent, which allows ML models to capture the recurring patterns in the dataset effectively. Minor errors were noted at the endpoints of certain predicted waveforms, primarily resulting from the inherent limitations of the initial segmentation process, where recordings are divided into segments of fixed durations.

Throughout the experiments, we notice that the consistency and uniformness of aphid’s behavior significantly affect the accuracy metrics used to assess the performance of the ML models, especially the OR metrics. Therefore, we present Fig 10, which demonstrates a situation in a recording from the SA dataset where the 2DCNN model attempts to annotate differently a long section of the first half of the recording in comparison to the ground-truth version. To be explicit, the upper panel displays the ground-truth annotation containing a long green section labeled as the C waveform. In the lower panel, different labels were given to this section, which are E1, E2, and NP (in light blue, pink, and light brown, respectively). In fact, the annotation by the ML model in this example is reasonable as we may observe behaviors resembling more the theoretical behavior of other feeding stages, especially the phloem-feeding stage.

**Fig 10.**
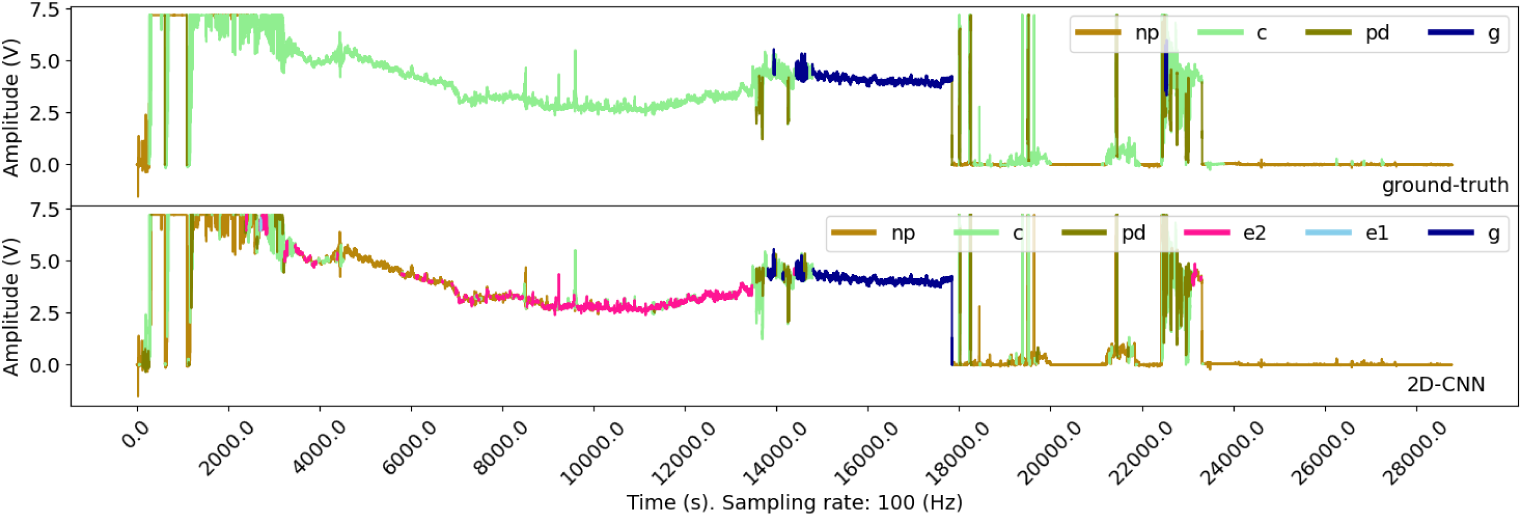
Example of Low Overlap Rate Due to Model Misclassification in SA Dataset. The figure presents an instance where the model’s predicted waveform annotation exhibits a low overlap rate with the ground-truth annotation in the SA dataset. Specifically, the misclassification of the model segment of the waveform is highlighted, demonstrating a potential area for model improvement or further investigation into the characteristics of the EPG signal in this specific dataset.

Although we experienced some false positive calls, ML models appear robust and consistent with the learnt pattern. The experimental results also support this claim, as we often observe the highest accuracy metrics in experiments where aphid behaviors experience minimal variation, such as in the BCOA1 and GPA datasets. In the BCOA2 and SA datasets, where the aphid’s behavior is not identical throughout the experiments, the predicted waveform often has a low overlap rate with the ground-truth version. Hence, it is essential to provide the ML models with well-annotated data for training to ensure the best performance.

Our study demonstrates several limitations that require further investigation. As waveform subsets extracted from a fixed-length *sliding-window* are treated *independently*, the models poorly annotate short waveforms such as pd that exist within C, a major waveform. The size of the sliding window also affects the level of offset between the endpoints of the predicted and ground-truth waveform, which we are willing to tolerate. A possible solution to this issue is to use more flexible region proposal algorithms such as Edge Boxes and Selective Search algorithms. These algorithms are well-known for object detection tasks, which propose potential regions containing objects (the EPG waveforms in our case) based on region information.

Removing the fixed boundaries restriction may allows the ML models to learn better the temporal structure of a waveform. In addition, it is possible to build an end-to-end DL model that can both learn these boundary positions and classify the waveform contained within. There exist deep learning models that were designed to handle similar problems in the image processing, such as those from the YOLO family or the R-CNN family, which can handle very well both the segmentation and detection tasks. For time-series adaptation, U-Time [26] is a CNN-based model that has shown effectiveness for sleep-staging, a similar problem to segmenting an EPG recording. We believe that by adopting these ideas, the reported results of our study can be greatly advanced.

Even while limitations may currently exist in characterizing EPG waveform, we have shown that the machine learning methods presented can achieve great accuracy and have the potential to greatly increase the productivity of EPG research. For example, the average EPG experiment can have a sample size ranging anywhere from 15-60 recordings, depending on the number of treatments. With the average recording taking anywhere from 10-30 minutes to annotate, this can lead to hours of annotation for a given experiment. Due to the cumbersome task of manual annotation, often this task is distributed amongst multiple researchers, which can lead to potential operator error and variation in the annotated waveforms. Thus, having accurate machine learning tools widely available to EPG researchers will allow for higher accuracy amongst waveform calls and allow for the users to increase sample sizes without risking the addition of large analysis timelines or high operator variability. In addition, an open-source Python package available to the public will enable EPG researchers to fine-tune and customize models that can be used to better fit their research needs. For example, a research group could create a custom model based on their labs recording data for a species of interest and pick and choose which models and model parameters work best in that particular system. The availability for EPG users to have access to machine learning tools allows for the field of EPG to gain the rigor associated with using computational tools rather than relying solely on the human eye. While further work needs to be done to allow even the most inexperienced users to utilize these tools, potentially via easily accessible web applications, this manuscript and the associated software packages provide a significant step in the introduction of machine learning tools for use in EPG waveform annotation. In conclusion, using machine learning tools in EPG research has the ability to enhance the accuracy of waveform classification across the field and significantly reduce the time and effort required for annotation.

## Supplementary Materials

**Fig S1.**
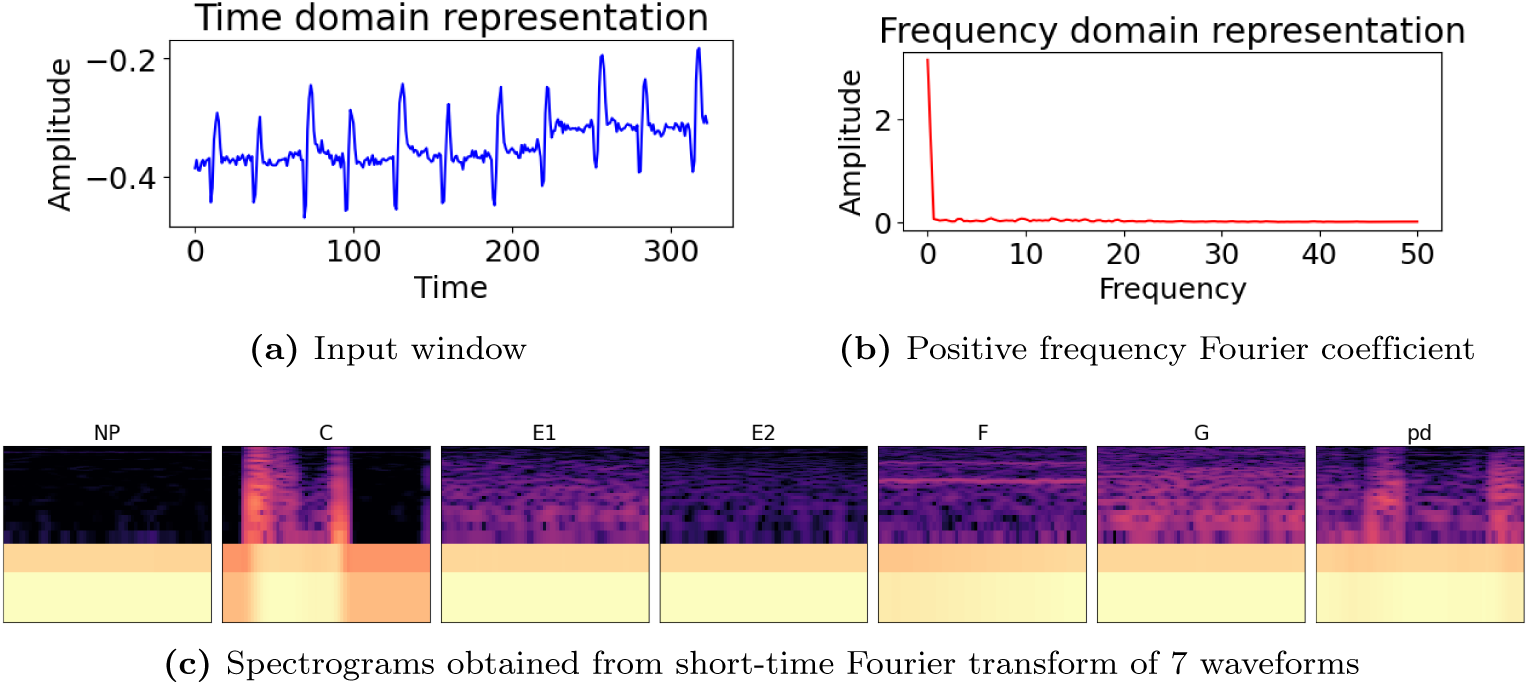
Fourier transform and spectrogram. The Fourier transform that converts a sliding window taken from E1 (a) into its amplitude-frequency representation (b) Studies have shown that waveforms can be identified based on the observed frequency, depending on the insect being studied. The spectrograms of the 7 waveforms are given in (c), which were calculated through short-time Fourier transform.

**Fig S2.**
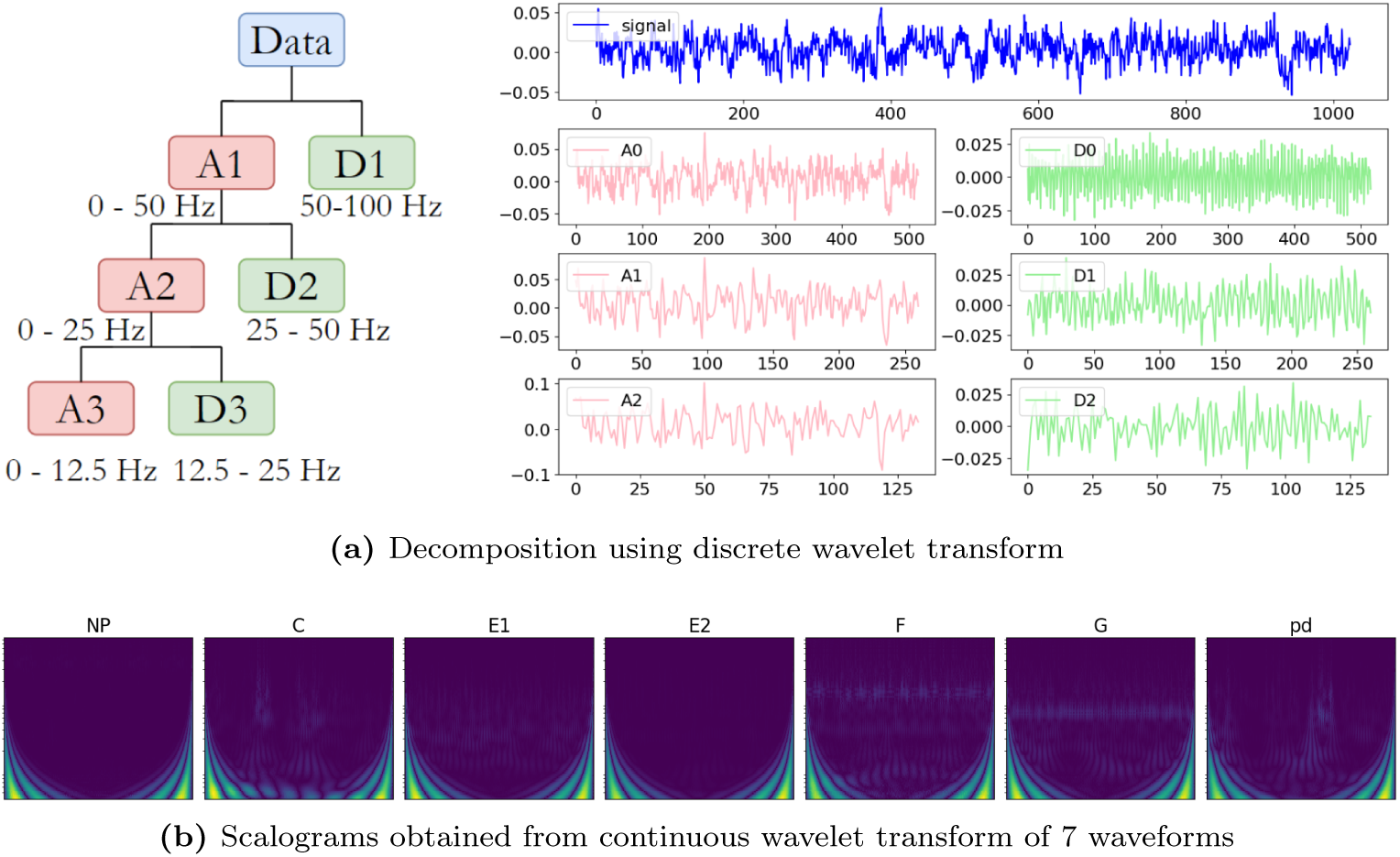
Wavelet transform and scalogram. The left panel displays the hierachical structure of the discrete wavelet transform decomposition. The entire signal is initially decomposed into approximation and detail coefficient, corresponding to low and high frequency, respectively. In each subsequent stage, the approximation coefficient are further decomposed in a similar manner. On the right hand side, an example of decomposing a fixed-length segment using wavelet transform with *symlet4* wavelet. Meanwhile, subfigure (c) presents the scalograms which were obtained using *Morlet* wavelet and continuous wavelet transform

**Fig S3.**
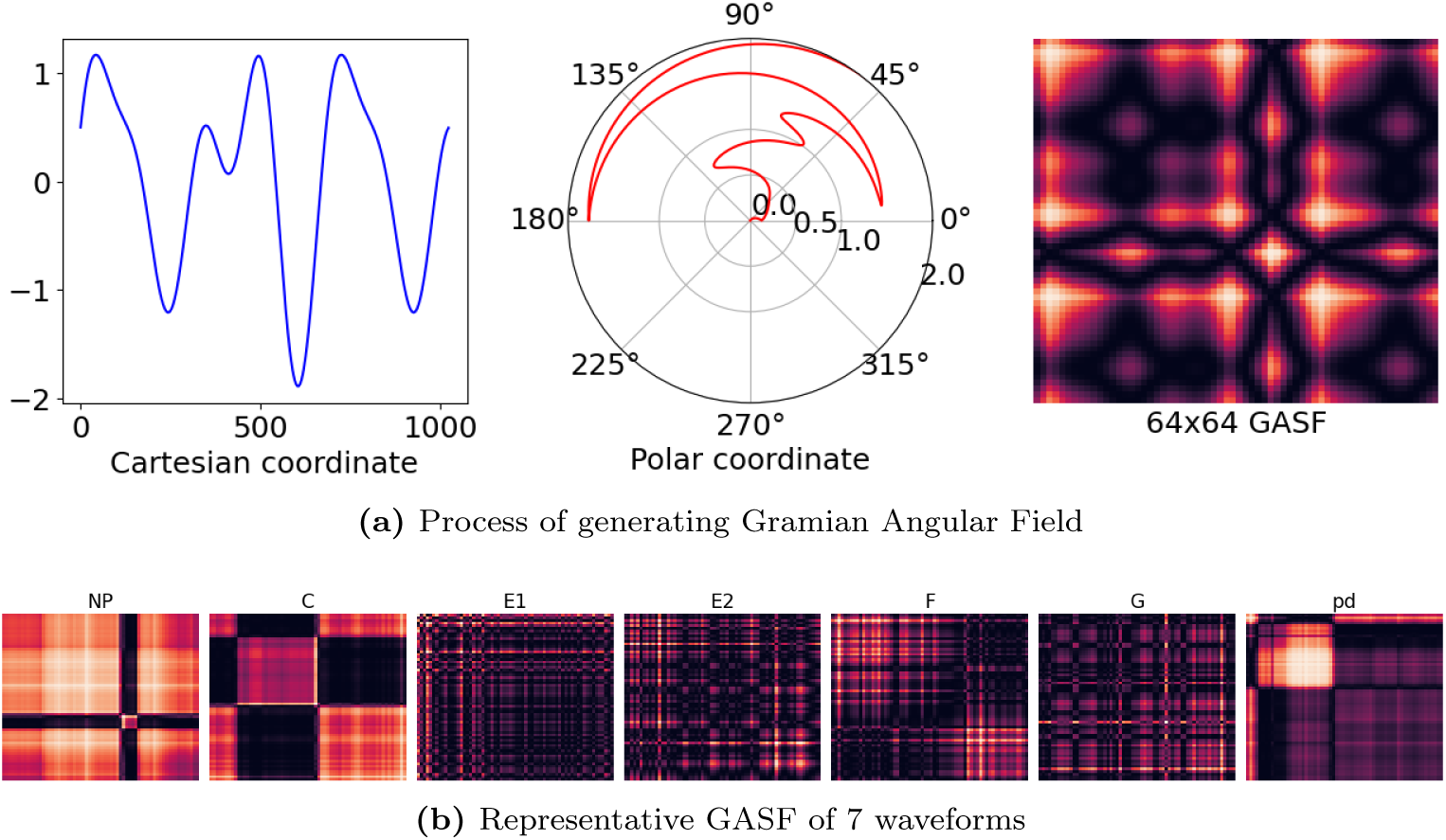
Gramian Angular Field. From left to right, subfigure (a) illustrates the process of generating a Gramian Angular Summation Field from an input time-series. The time-series are transformed into polar coordinate, then convert into a matrix consisting of trigonometrics values between pairs of angles, which represents an image. In subfigure (b), we shows the represntative GASF of 7 waveforms.

**Fig S4.**
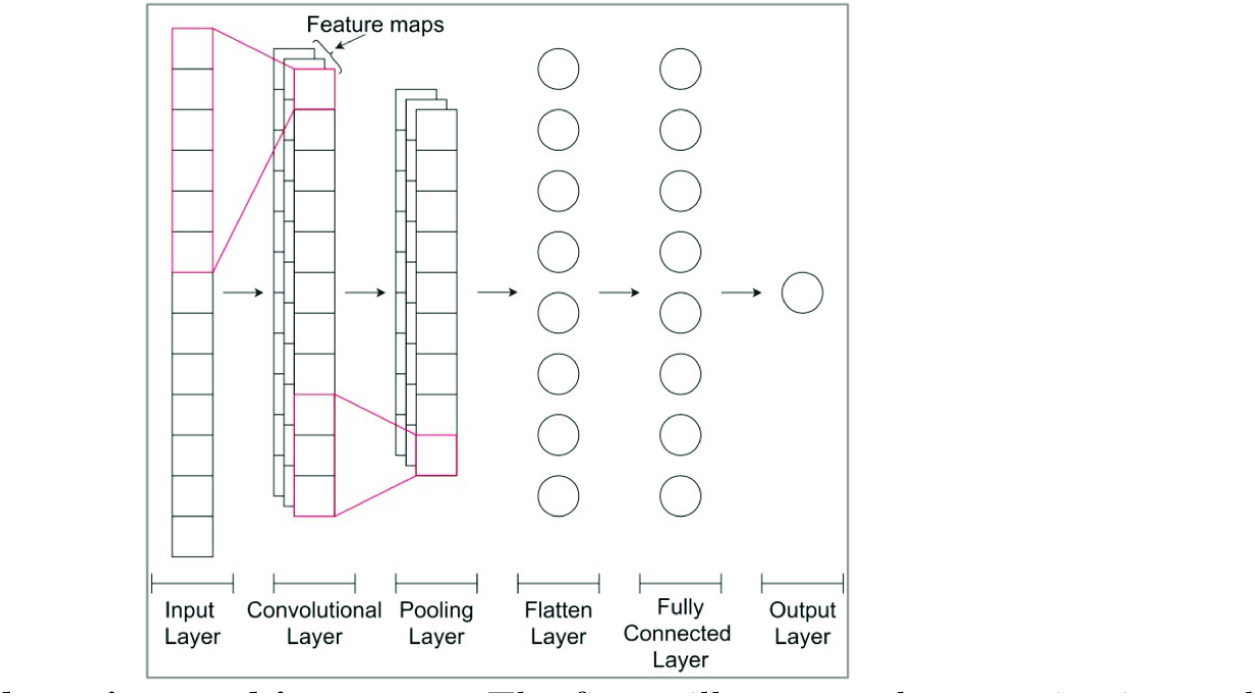
Deep learning architectures. The figure illustrates the organization and the types of layers founded in a Convolutional Neural Network (CNN).

